# A *Drosophila in vivo* chemical screen reveals that combination drug treatment targeting MEK and DGKα mitigates Ras-driven polarity-impaired tumourigenesis

**DOI:** 10.1101/2022.03.14.484232

**Authors:** John E. La Marca, Robert W. Ely, Sarah T. Diepstraten, Peter Burke, Gemma L. Kelly, Patrick O. Humbert, Helena E. Richardson

## Abstract

The *RAS* oncogene and upregulation of the RAS signalling pathway is highly prevalent in human cancer, and therefore, therapeutically targeting the RAS pathway is a common treatment in cancer. However, RAS pathway upregulation is not sufficient to drive malignant cancer, since senescence mechanisms prevent cancer progression. Thus, additional mutations, such as mutations that prevent senescence or alter the tissue architecture (cell polarity), are required for *RAS*-driven tumour progression. Moreover, targeting *RAS*-driven cancers with RAS pathway inhibitors can often lead to undesirable side-effects and to drug resistance. Thus, identifying compounds that synergise with RAS-pathway inhibitors would enable lower doses of the RAS pathway inhibitors to be used and also decrease the acquisition of drug resistance. Here, in a boutique chemical screen using a *Drosophila* model of Ras-driven cell polarity-impaired cancer, we have identified compounds that reduce tumour burden by synergising with subtherapeutic doses of the RAS pathway inhibitor, Trametinib, which inhibits mitogen-activated kinase kinase (MEK). Analysis of one of the hits from the screen, Ritanserin, which targets serotonin receptors and diacy glycerol kinase alpha (DGK*α*), revealed that DGK*α* was the critical target in its synergism with Trametinib. We show that human mammary epithelial cells harbouring the *H-RAS* oncogene and knockdown of the cell polarity gene, *SCRIB*, are also sensitive to treatment with low doses of Trametinib and DGK*α* inhibition. Mechanistically, DGK*α* inhibition synergises with Trametinib by inhibiting MEK and mTOR activity. Altogether, our results provide evidence that targeting RAS-driven human cancers with RAS pathway and DGK*α* inhibitors will be an effective combination therapy.

## Introduction

Cancer places a huge burden on public health outcomes worldwide, with estimated new cancer cases numbering over 14 million annually, and associated deaths approximated at 8.2 million for the year 2012 (Ferlay et al., 2015; Siegel et al., 2016). Coupled with these staggering rates of incidence are the difficulties involved in creating new and improved treatments for cancer. Emerging anti-cancer drugs have the lowest likelihood of moving forward from phase 1 trials compared to all other classes of drugs, and the list of new drug candidates gaining US Food and Drug Administration (FDA) approval for clinical use leaves much to be desired (Hay et al., 2014). Thus, the field of oncology could benefit from enhanced and more immediate methods for drug discovery, or the repurposing of already FDA approved drugs. One model that can aid in this process is the vinegar fly, *Drosophila melanogaster*, which can be used to screen for novel anti-cancer compounds *in vivo*.

*Drosophila* is used extensively in the study of many genetic diseases (Bier, 2005; Wangler et al., 2015; Wangler et al., 2017), and has been used for the investigation of cancer for over 100 years (Gonzalez, 2013; Stark, 1918). *Drosophila* carries orthologues of 68% of known human cancer-causing genes, and there is also a high level of conservation between human and *Drosophila* biological processes and signalling pathways (Brumby and Richardson, 2005; Hanahan and Weinberg, 2011). These factors, as well as its short life-cycle and low maintenance costs, positions *Drosophila* as an invaluable tool, filling a niche between mammalian cell lines and more complex organisms for the study of cancer both *in vitro* and *in vivo,* and also enables its use as a platform for the identification of anti-cancer compounds (Bangi, 2019; Brumby and Richardson, 2005; Gladstone and Su, 2011; Gonzalez, 2013; Richardson et al., 2015; Rudrapatna et al., 2012). Importantly, a majority of the “hallmarks of cancer” can be modelled in *Drosophila*, including increased cell proliferation, evading apoptosis and differentiation, and induction of invasion/metastasis (Brumby and Richardson, 2005; Hanahan and Weinberg, 2011).

A group of genes heavily implicated in human cancers are the oncogenic *RAS* genes, which signal through the RAF-MEK-MAPK and Phospho-inositol-3-kinase (PI3K)-AKT-mTOR pathways to drive cell growth, proliferation, and survival (Malumbres and Barbacid, 2003; Pylayeva-Gupta et al., 2011). However, oncogenic mutations in *RAS* are not sufficient to drive malignant cancers since high levels of RAS signalling leads to cell-cycle arrest and senescence, and therefore additional mutations are needed to overcome these curbs to cancer progression (Coleman et al., 2006; DeNicola and Tuveson, 2009; Dimauro and David, 2010; Olson et al., 1998; Sahai et al., 2001). The *Drosophila* orthologue of the human *RAS* genes is *Ras85D* (hereafter termed *Ras*). Expression of an activated form of *Drosophila Ras*, termed *Ras^V12^* (which bears the constitutively activating G12V mutation), induces hyperplastic growth in a variety of tissues, but further tumour progression does not occur due to cell cycle arrest, differentiation and senescence-like characteristics (Brumby et al., 2011; Ito and Igaki, 2021; Karim and Rubin, 1998; Nakamura and Igaki, 2017).

A major factor in tumourigenesis is the loss of cell polarity, with the majority of human epithelial tumours estimated to have cell polarity and tissue architecture disruption (Godde et al., 2014; Lee and Vasioukhin, 2008; Muthuswamy and Xue, 2012; Royer and Lu, 2011). Indeed, the disruption of cell polarity is implicated as a causative factor in many different cancers (such as breast and cervical cancers (Feigin et al., 2014; Thomas et al., 2008)), with cell polarity genes acting as so-called “tumour-suppressors” (Bilder, 2004; Sonoshita and Cagan, 2017). *Drosophila lethal (2) giant larvae* (*l(2)gl*), *discs large 1* (*dlg1*), and *scribble* (*scrib*) are cell polarity genes that, when mutated, result in cells exhibiting a loss of polarity and tissue architecture, disrupted differentiation, and increased tissue growth. Upon transplantation into adult flies, these mutant cells massively overgrow and undergo invasion/metastasis reminiscent of mammalian cancers (Froldi et al., 2008). Mammalian orthologs of L(2)gl, Dlg1 and Scrib similarly act as tumour suppressors, restraining cell proliferation and invasion/metastasis (Elsum et al., 2012; Humbert et al., 2008; Stephens et al., 2018).

When *Drosophila* cell polarity genes are mutated, the tissue exhibits overgrowth through impairment of the Hippo pathway, a negative tissue growth control pathway whose downstream target, Yorkie (Yki), functions as a co-transcriptional activator to drive expression of the cell growth/proliferation genes, *myc* and *Cyclin E*, and the anti-apoptotic gene, *Death-associated inhibitor of apoptosis 1* (*Diap1*), thereby causing increased cell proliferation and survival (Doggett et al., 2011). When disrupted in clones within a whole tissue, mutant *scrib* cells do not display the phenotype of aggressive tumours and are largely eliminated via c-Jun N-terminal Kinase (JNK)-mediated apoptosis (Brumby and Richardson, 2003; Leong et al., 2009). However, when *Ras^V12^* is expressed within epithelial tissues that are also mutant for *scrib*, neoplastic tumours are generated, which show increased cell proliferation, increased survival, reduced differentiation, and exhibit invasive/metastatic behaviour, thereby replicating many of the mammalian cancer hallmarks (Brumby and Richardson, 2003; Leong et al., 2009; Pagliarini and Xu, 2003). Alone, *Ras^V12^* drives cell proliferation and survival, but also induces cell cycle arrest, senescence and differentiation (Brumby et al., 2011; Ito and Igaki, 2021; Karim and Rubin, 1998; Nakamura and Igaki, 2017). Aggressive and neoplastic properties arise through the cooperative effects of these two mutations, largely through combination of the tissue overgrowth consequences and suppression of senescence by impaired Hippo pathway signalling, the pro-survival characteristics induced by Ras activation, and the hijacking of activated JNK signalling to block differentiation and induce invasion through the upregulation of matrix metalloproteases (MMPs) (Doggett et al., 2011; Igaki et al., 2006; Ito and Igaki, 2021; Leong et al., 2009; Uhlirova and Bohmann, 2006).

The simplified genetics of the fly (with fewer redundant genes than in mammals allowing for easier knock-down models), together with the ability to rear hundreds of animals that can easily have their diet supplemented with drugs and monitored in a whole-body environment, are powerful benefits to using *Drosophila* in a drug discovery setting (Bangi, 2019; Gladstone and Su, 2011; Richardson et al., 2015; Yadav et al., 2016). An analysis of drugs specifically targeting various signalling pathways in *Drosophila* revealed that the mode of action of most was conserved between flies and humans (Bangi et al., 2011), reinforcing the utility of *Drosophila* as a platform for drug discovery that is relevant to human biology. Indeed, several studies have utilized *Drosophila* as a model system for anti-cancer drug discovery, using a variety of approaches. One of the earliest examples used the suppression of a *Drosophila* eye phenotype caused by the expression of the *Ret* oncogene to validate the compound ZD6474 (a small molecule receptor tyrosine kinase inhibitor, known now as Vandetanib) as a potential therapy for *Ret* oncogene-driven medullary thyroid cancer (MTC), a disorder observed in multiple endocrine neoplasia type 2B (MEN2B) (Vidal et al., 2005). A similar approach has been used to test further compounds for efficacy against *Ret*-induced phenotypes, identifying the compound AD57, which was then validated in human cell lines (Dar et al., 2012). The continued refinement of this model has led to the development of ‘tumour calibrated inhibitors’, where using genetic and chemical modifier screens, existing kinase inhibitors can be modified to create new compounds with more precise targeting and superior efficacy (Masahiro et al., 2018).

Large-scale chemical screens have also been employed in the *Drosophila* adult intestine to identify compounds that target RAF-driven tumours (Markstein et al., 2014), as well as to reveal compounds that target various molecularly-defined types of human colorectal cancer (Bangi et al., 2019; Bangi et al., 2016). Similarly, using the adult *Drosophila* trachea (breathing tubes) as a model for human lungs, large-scale chemical screens have identified compounds that target *RAS*-driven lung cancers (Levine and Cagan, 2016). *Drosophila* as a large-scale screening platform has also been effectively used in the search for drugs that intensify the effects of radiation *in vivo* by identifying compounds that reduced survival of *Drosophila* larvae after irradiation (Edwards et al., 2011; Gladstone et al., 2012; Jaklevic et al., 2006). In this manner, the translational elongation inhibitor Bouvardin was identified to act in a synergistic manner with irradiation to reduce organismal survival, a result which was then validated in mammalian cells lines and human cancer xenografts in mice (Gladstone et al., 2012).

In our previous studies, we utilized a clonally-induced, polarity-impaired Ras-driven (*scrib^-^/ Ras^V12^* model of cancer to screen a 2000 compound library and identify drugs that reduced tumour burden in *Drosophila* larvae (Willoughby et al., 2013). We developed a screening platform in a 96-well plate format, where larvae were fed food containing different compounds for 5 days, before being imaged to assess the effect on GFP-marked tumour size (Willoughby et al., 2013). In this screen we identified Acivicin, a glutamine analogue already known to have anti-tumour properties in humans, and upon pharmacogenetic analysis demonstrated that it targeted the glutamine-utilization enzyme CTP synthase, as well as the TCA cycle (Willoughby et al., 2013).

In this current study, we adapt our *scrib^-^/Ras^V12^* model and the larval screening platform we developed to screen for compounds that synergistically inhibit tumourigenesis together with the MEK (mitogen activated protein kinase kinase) inhibitor, Trametinib. Trametinib is an orally-administered, FDA-approved, widely used drug for the treatment of Ras-driven cancers, including melanoma, non-small cell lung cancer, thyroid cancer, and glioma (Ferrari et al., 2020; Hoffner and Benchich, 2018; Manoharan et al., 2020; Tabbo et al., 2022; Wright and McCormack, 2013; Zeiser, 2014). However, Trametinib has dermatological side-effects that often lead to the dose being lowered or treatment withdrawn (Abdel-Rahman et al., 2015; Anforth et al., 2014). Thus, identifying compounds that can effectively synergize with Trametinib to inhibit Ras pathway-driven cancers, enabling lower doses of Trametinib to be used, would be highly desirable. After screening ∼5000 compounds, we identified 2 compounds that synergized with a sub-therapeutic dose of Trametinib to reduce tumour burden: the Polo-like kinase inhibitor, Volasertib, and the serotonin receptor and diacy glycerol kinase (DGK) inhibitor, Ritanserin. Further phamacogenetic analyses of the mode of action of Ritanserin indicated that inhibition of DGK is required to reduce tumour burden in cooperation with Trametinib. We demonstrate that this synergistic mechanism is conserved in human mammary epithelial cells, with the drug combination specifically targeting Ras-driven, polarity-impaired cells with minimal effects on normal cells. Thus, low dose Trametinib combined with Ritanserin, or with more selective DGK inhibitors, is a novel anti-cancer combination therapy that could be developed to target human mammary cancers, as well as other human Ras-driven polarity-impaired cancers.

## Results

### Identification of compounds that synergise with a sub-therapeutic dose of the MEK inhibitor, Trametinib, to suppress polarity-impaired Ras-driven tumours in *Drosophila*

Using the clonally-induced *scrib* mutant, oncogenic *Ras* (*Ras^V12^*) model of epithelial tumourigenesis (Brumby and Richardson, 2003; Leong et al., 2009) we have previously shown that feeding tumour-bearing larvae a bioavailable compound (PD0325901) that targets MEK (*Drosophila* Dsor1), a protein kinase in the Ras-MAPK pathway, was effective in reducing tumour burden (Willoughby et al., 2013). Since another MEK inhibitor, Trametinib, is a widely used orally-administered drug for the treatment of Ras-driven human cancers (Ferrari et al., 2020; Hoffner and Benchich, 2018; Manoharan et al., 2020; Tabbo et al., 2022; Wright and McCormack, 2013; Zeiser, 2014), we tested if it was also effective in reducing tumour burden of *scrib* mutant *Ras^V12^*-expressing (*scrib^-^*/*Ras^V12^*) larvae when administered in the food. Indeed, we found that Trametinib was highly effective in reducing the tumour size in these tumour-bearing larvae (Fig 1). We then titrated the concentration of Trametinib to determine the minimal dose that resulted in a significant, visible reduction of GFP-marked tumour burden, and found that a final concentration of 2.5*μ*M Trametinib led to a 70-50% reduction of tumour size (Fig 1). Thus, we hypothesized we would be able to identify compounds that could synergize with Trametinib in inhibiting tumour burden by using the sub-therapeutic dose of 2.5*μ*M Trametinib in a chemical screen.

**Figure 1.**
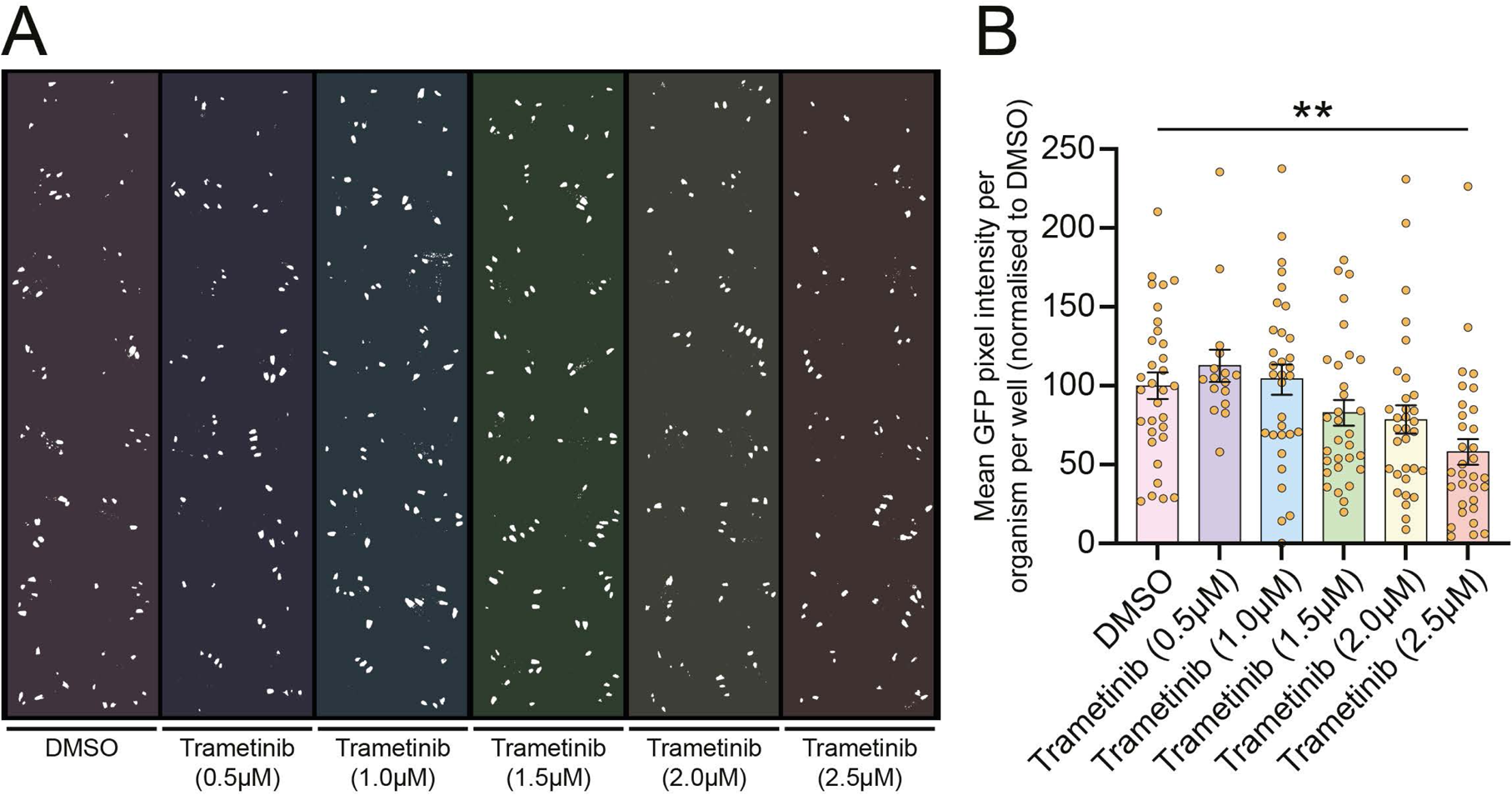
Titration of Trametinib fed to *scrib^-^/Ras^V12^* tumour-bearing larvae reveals the minimum concentration needed for tumour growth inhibition. (A) To determine the optimal concentration of the MEK inhibitor, Trametinib, for synergism assays, we titrated the drug across a plate of larvae possessing *scrib^-^/Ras^V12^* tumours. (B) Quantifications of GFP pixel intensity revealed that, at a final concentration of 1.5µM and above, Trametinib reduced the tumour burden in the larvae, achieving knockdown to ∼60% of the DMSO-treated control. Results represent 2 replicate experiments with 16 (for Trametinib at 0.5µM) or 32 (for each other treatment) replicate wells for each treatment. Statistical test used was a one-way ANOVA with Tukey’s multiple comparisons, with non-significant comparisons not visualised. Error bars represent S.E.M. ** = p<0.01.

To identify compounds that could synergise with a Ras-MAPK pathway inhibitor, Trametinib, we screened 4 boutique compound libraries (obtained from the Walter and Eliza Hall Institute): kinase inhibitors (276 compounds), epigenetic modifiers (89), targeted compounds (179), known drugs (3707) (File S1). We assessed their effectiveness at reducing *scrib^-^/Ras^V12^* tumour burden combined with a sub-therapeutic dose of Trametinib (2.5*μ*M), which only minimally affected the tumour burden in isolation. We screened the tumour-bearing larvae in 96 well deep-well micro-titre plates where different drugs were administered in the food at a final concentration of 50*μ*M, along with DMSO control wells, as previously described (Richardson et al., 2015; Willoughby et al., 2013). We screened the first 3 libraries in duplicate +/- sub-therapeutic levels of Trametinib, but since the known drug library was in limited supply, we only screened one copy of this library in the presence of Trametinib. From this screen we identified 20 potential hits that reduced tumour burden in all larvae in the well relative to the DMSO control wells for each plate, but which did not appear to reduce overall larval size. These compounds included Trametinib itself, as well as compounds targeting a variety of other pathways (Table 1). To validate the candidates, we tested independently-sourced supplies of these 19 novel compounds, +/- a sub-therapeutic dose of Trametinib in multiple wells of the micro-titre plates. This analysis revealed that 13 of the compounds did not significantly reduce the size of *scrib^-^/Ras^V12^* tumours (File S2). Four compounds, Methotrexate and Pralatrexate (folate antagonists), Temsirolimus (mTOR inhibitor), and GSK2126458 (PI3K inhibitor), were confirmed to reduce tumour burden, but did not synergize with Trametinib at the various doses tested (Figs S1 and S2), and therefore were not further analyzed. Importantly, two compounds were confirmed to display synergism with Trametinib in reducing tumour burden: Volasertib (Polo-like kinase inhibitor (Schoffski, 2009)) and Ritanserin (Serotonin Receptor 5-HT2A/2C and diacyl glycerol kinase α (DGKα) inhibitor (Boroda et al., 2017; Leysen et al., 1985)) (Fig 2). At 2 different doses, Volasertib showed a reduction in tumour burden on its own, but at the lower dose (12.5*μ*M) showed synergism with Trametinib (Fig 2A, B). Ritanserin (50*μ*M) did not show any significant effect on tumour burden on its own but synergised with Trametinib to significantly reduce tumour burden from ∼75% to 50% (Fig 2C, D). However, at lower doses of Trametinib (1.25*μ*M), Ritanserin (50*μ*M) was unable to reduce tumour size (Fig S3), indicating that at least 2.5*μ*M Trametinib is required for a robust synergistic response with Ritanserin. Thus, we have discovered two compounds, Volasertib (Polo-like kinase inhibitor) and Ritanserin (Serotonin Receptor 5-HT2A/2C and DGKα inhibitor) that synergise with Trametinib to reduce *scrib^-^/Ras^V12^* tumour burden. Since the Polo-like kinase inhibitor, Volasertib, had an effect on its own, and as this class of inhibitors have already been extensively utilised in cancer therapy (Gjertsen and Schöffski, 2014; Shakeel et al., 2021; Zhang et al., 2021), we focused our attention on Ritanserin, which had no significant effect on tumour burden on its own, and has no history of use as an anti-cancer drug.

**Figure 2.**
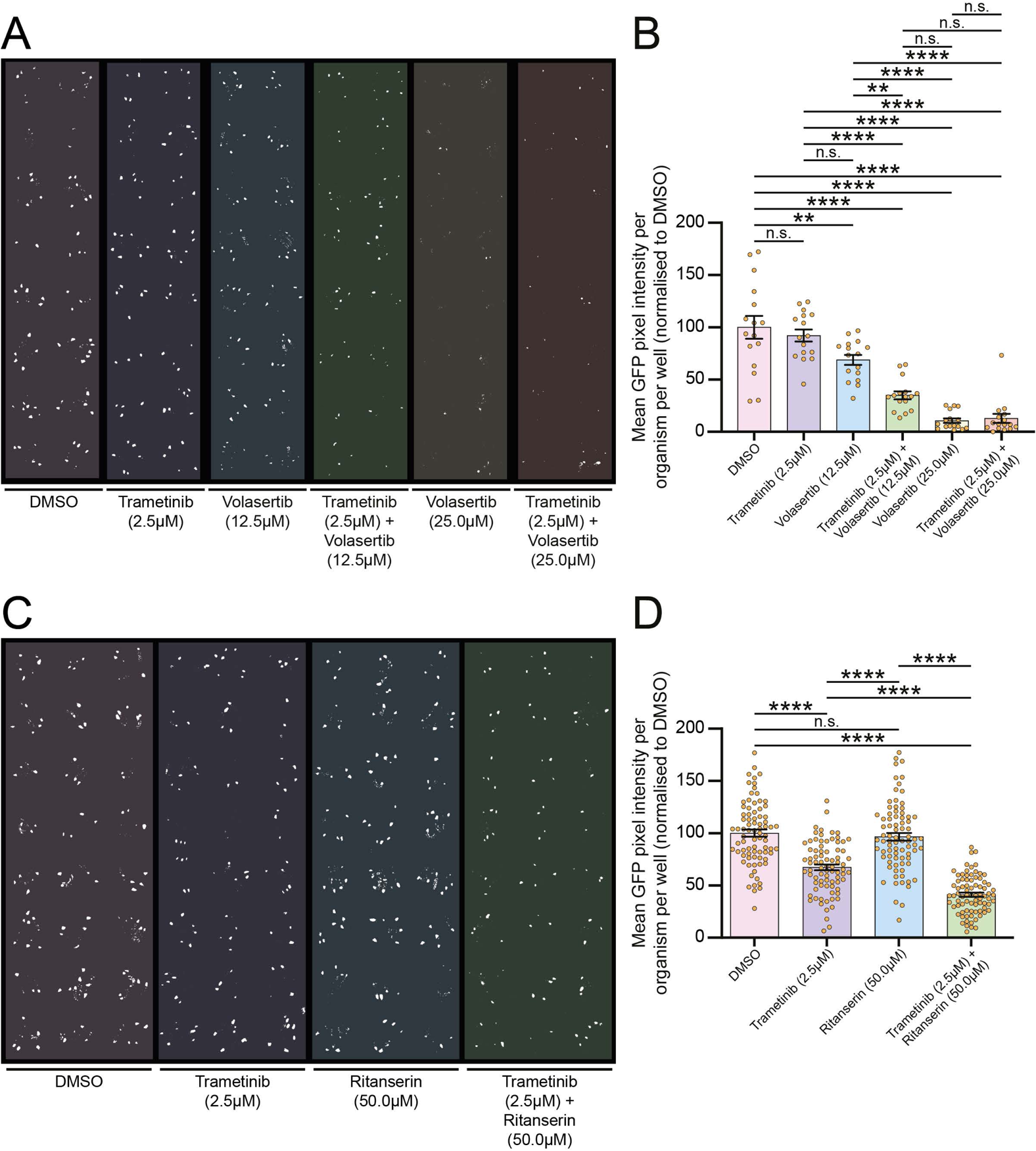
Volasertib and Ritanserin show synergy with Trametinib in reducing the tumour burden in *scrib^-^/Ras^V12^* tumour-bearing larvae. (A) Binarized image of a drug test plate where larvae were treated with Trametinib in combination with the Polo-like kinase inhibitor, Volasertib. (B) Quantifications of GFP pixel intensity revealed that Volasertib synergised with Trametinib at final concentrations of 12.5µM and 2.5µM, respectively, to significantly reduce tumour burden to ∼35% of the DMSO-treated control. However, increasing the Volasertib concentration to 25µM led to tumour-burden decrease independent of Trametinib. Data from 1 experiment with 16 replicate wells for each treatment. (C) Binarized image of a drug test plate where larvae were treated with Trametinib in combination with the serotonin receptor and diacyl glycerol kinase inhibitor, Ritanserin. (D) Quantifications of GFP pixel intensity revealed that Ritanserin synergised with Trametinib at final concentrations of 50µM and 2.5µM, respectively, to significantly reduce tumour burden to ∼40% of the DMSO-treated control. Data from 4 replicate experiments with 16-24 replicate replicate wells for each treatment. Statistical tests used were one-way ANOVAs with Tukey’s multiple comparisons. Error bars represent S.E.M. ** = p<0.01, *** = p<0.001, **** = p<0.0001.

**Table 1.**
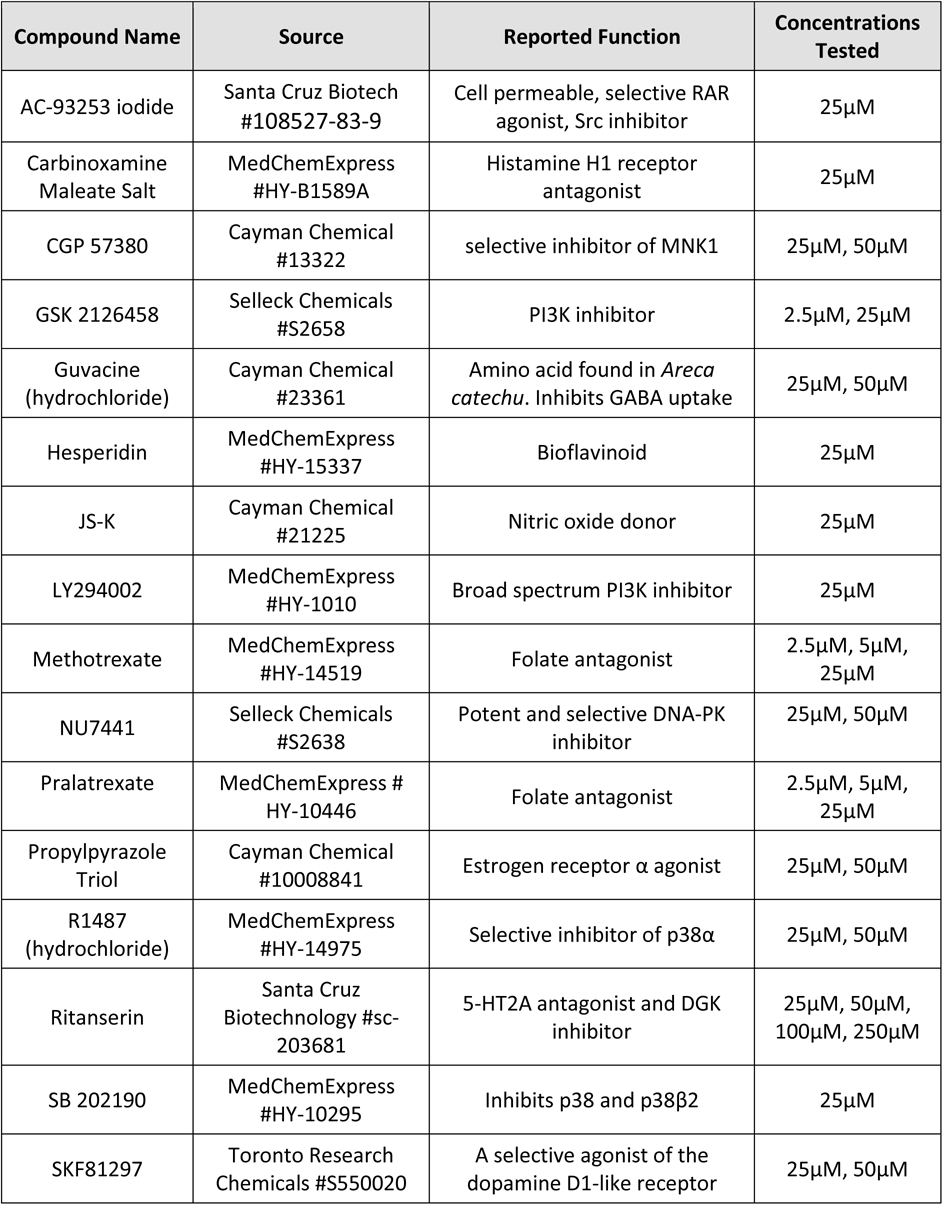
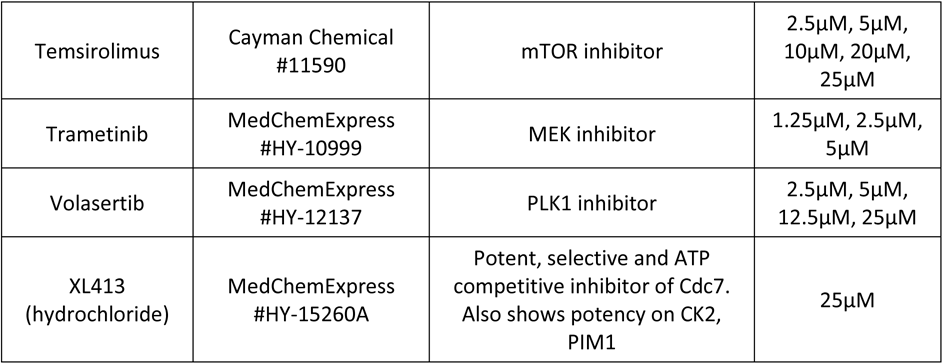
Compounds potentially synergistic with Trametinib. This table lists compounds identified in the primary screen that were potentially synergistic with Trametinib at reducing *scrib^-^/Ras^V12^* tumour burden. Follow-up experiments at the concentrations listed tested this possibility of synergy.

### Ritanserin synergises with Trametinib to target tumour growth and enable continued development of *scrib^-^/Ras^V12^* tumour-bearing *Drosophila* larvae

First, we sought to determine whether Ritanserin +/- Trametinib was specifically targeting the *scrib^-^/Ras^V12^* tumour, rather than affecting growth of the whole organism. To do this, we measured the size of both whole larvae and their respective GFP-labelled tumours (n=78 per treatment) in drug-treated or DMSO vehicle treated control samples (Fig 3). We found that Ritanserin treatment alone resulted in a small increase in the average larval size (Fig 3A). By contrast, the low dose of Trametinib did not affect *scrib^-^/Ras^V12^* tumour-bearing larval size, whilst the combined treatment of Trametinib with Ritanserin resulted in a small decrease in the average larval size (Fig 3A). However, when the effect of the drugs on GFP-marked tumour size was presented as a ratio compared with whole larval size, Ritanserin alone had no effect, Trametinib reduced the tumour to ∼60% of the DMSO control, and the combined drug combination showed a reduction of the relative tumour size to ∼35% of the DMSO control (Fig 3B). This result indicates that the Ritanserin + Trametinib drug combination is targeting the tumour specifically.

**Figure 3.**
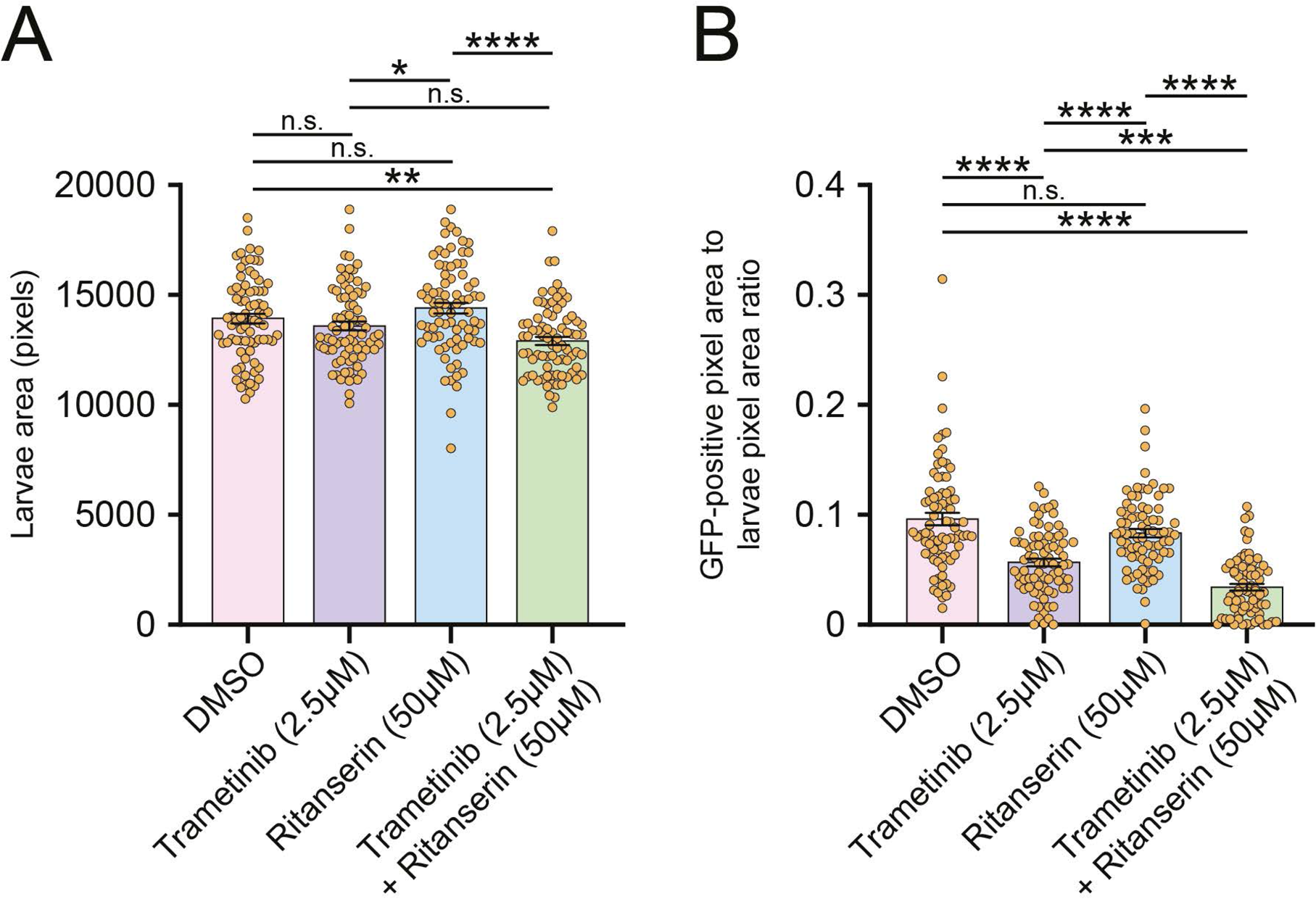
Trametinib and Ritanserin combination treatment directly targets *scrib^-^/Ras^V12^* tumour growth. (A) Quantifications of larval size after treatment with Trametinib and Ritanserin revealed that there was a significant reduction in larval size in the larvae undergoing combination treatment compared to the DMSO control, but there was no difference in larval size whether treated with Trametinib alone or Trametinib and Ritanserin in combination. (B) Quantification of the ratio between GFP pixel intensity and larval area demonstrate that despite the reduction in larval size, Trametinib and Ritanserin combination treatment leads to a tumour size of ∼3.5% of the larvae area, a significant reduction compared to ∼9% in the DMSO control and Ritanserin-alone treatments, and ∼5.5% in the Trametinib-alone treatment. Data derived from 2 replicate experiments with 16-23 replicate wells for each treatment. Statistical test used was a one-way ANOVA with Tukey’s multiple comparisons. Error bars represent S.E.M. * = p<0.05, ** = p<0.01, *** = p<0.001, **** = p<0.0001.

We next examined whether the Ritanserin +/- Trametinib drug treatment could rescue development of the *scrib^-^/Ras^V12^* tumour-bearing larvae, which generally are overgrown due to a block or delay in progressing to the pupal stage. To test this, we fed drugs to the developing larvae via their food in standard *Drosophila* vials (transferred at the 2^nd^ instar larval stage) and observed the number of pupae that developed (Fig 4). In this scaled-up experiment, conducted on molasses-rich fly food, higher doses of the drugs (final concentrations of Trametinib at 10*μ*M and Ritanserin at 200*μ*M) were needed to produce effects. We found that whilst only ∼5% of DMSO-treated *scrib^-^/Ras^V12^* tumour-bearing larvae pupated, and ∼40% of Trametinib-treated larvae produced pupae, the combination of Ritanserin + Trametinib resulted in 90% of the larvae pupating (Fig 4E). Furthermore, whilst many of the larvae from the DMSO or Trametinib treated samples died before reaching the 3^rd^ instar larval stage, the drug combination dramatically increased organism survival relative to DMSO or Trametinib alone (Fig 4E). Indeed, the Ritanserin + Trametinib combination allowed the formation of adult body structures, including the head and wings, which were visible through the pupal case (Fig 4D, compared with the *wild-type* control in Fig 4A, and DMSO treated tumour-bearing larvae in Fig 4B), whilst Trametinib alone only allowed the development of head structures in some pupae (Fig 4C). However, although the combination drug treatment allowed development to occur in the pupal stage, none of these pupae were able to eclose and produce adult flies. Thus, the Ritanserin + Trametinib drug combination increases the survival of tumour-bearing larvae and increases their ability to progress into the pupal stage and form adult structures, which is consistent with the effect of the drug combination in reducing tumour size.

**Figure 4.**
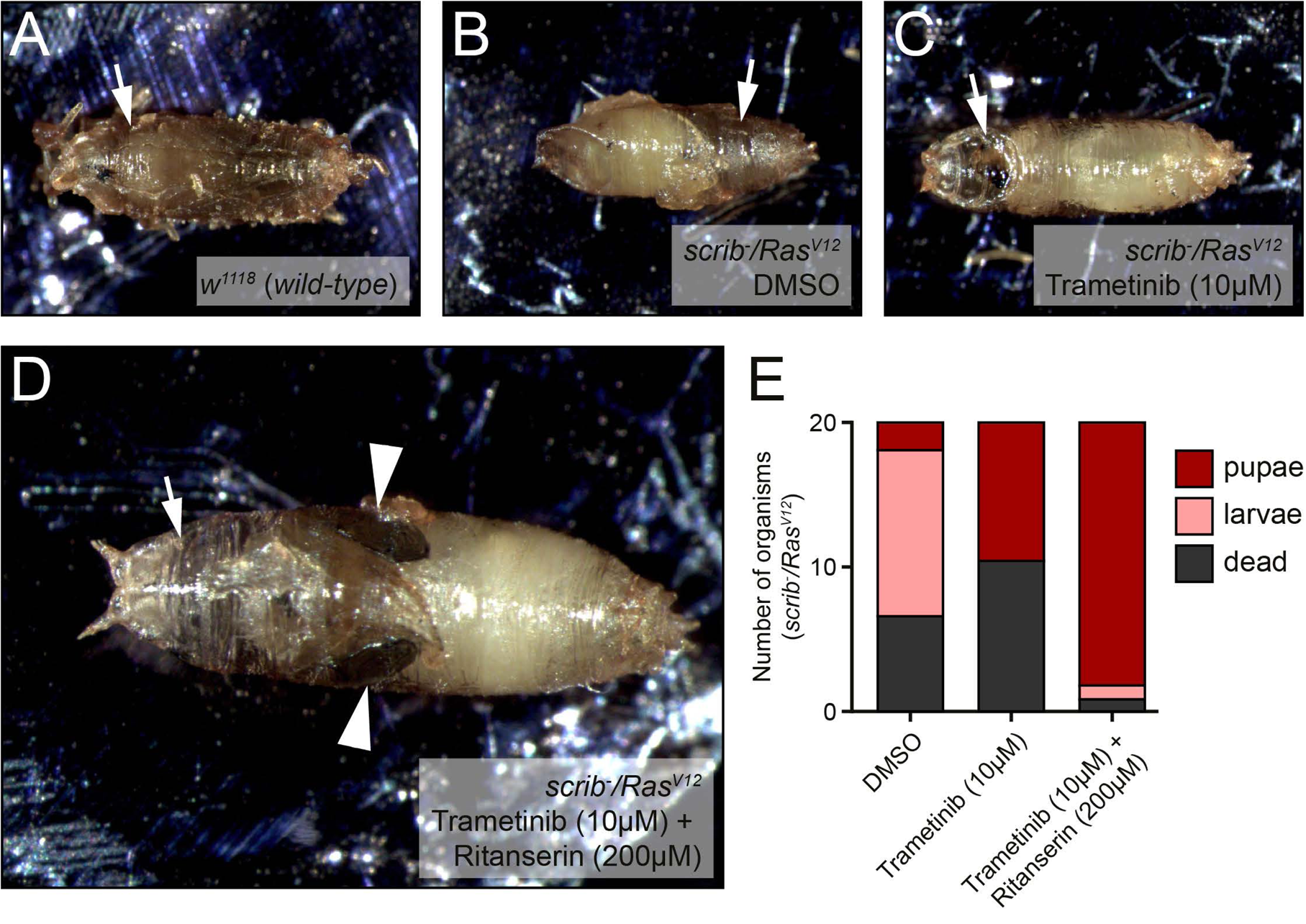
Trametinib and Ritanserin combination treatment partially rescues pupal development of *scrib^-^/Ras^V12^* tumour-bearing animals. (A) 7-8 day old *wild-type* (*w^1118^*) pupae demonstrates regular development, with proper formation of the adult head and body (arrow). (B) Lethality of the *scrib^-^/Ras^V12^* genotype is apparent at the pupal stage, with improper metamorphosis and body formation (arrow) observable. (C) Treatment of *scrib^-^/Ras^V12^* animals with Trametinib (10µM, 10 days) results in more pupae, but reduced head development (arrow) and no clear adult structures being observed. (D) Treatment of *scrib^-^/Ras^V12^* animals with Trametinib and Ritanserin (10µM and 200µM, respectively, 10 days) results in sufficient rescue of development that adult wing structures could be observed (arrowheads), though head development was still incomplete (arrow). (E) Quantification of the *scrib^-^/Ras^V12^* pupae. Trametinib and Ritanserin combination treatment resulted in increased pupation, but eclosion was never observed. For each treatment, n=21 animals.

### Ritanserin targets Dgk to synergise with Trametinib in inhibiting Ras-driven polarity-impaired tumourigenesis in *Drosophila*

Since Ritanserin inhibits both DGKα and serotonin receptors (Boroda et al., 2017; Franks et al., 2017; Mizutani et al., 2018), we wished to determine whether Dgk or serotonin receptors are the key target for Ritanserin in *Drosophila* in reducing *scrib^-^/Ras^V12^* tumour burden in cooperation with Trametinib by analysing specific DGK*α* or serotonin receptor inhibitors. Firstly, to confirm the results obtained with Ritanserin, we analysed another DGKα/serotonin receptor inhibitory drug, R-59-022 (Boroda et al., 2017; Sato et al., 2013). Treatment of tumour-bearing larvae with R-59-022 together with a low dose of Trametinib resulted in a significant reduction in tumour size (Fig S4A,B) but had no significant effect alone, thereby complementing the results with Ritanserin. We then analysed the DGKα specific inhibitors, Amb639752 (Velnati et al., 2019) and CU-3 (Liu et al., 2016; Yamaki et al., 2019). When treated with the DGKα-specific inhibitors, Amb639752 or CU-3, together with a low dose of Trametinib, significant decreases in tumour burden were also observed (Fig S4C-F), indicating that Dgk is the relevant target of Ritanserin in its cooperation with Trametinib in inhibiting Ras-driven polarity-impaired tumourigenesis. Consistent with these findings, treatment of the tumour-bearing larvae with specific serotonin receptor inhibitors, Volinanserin (which has high selectivity for the 5-HT2A serotonin receptor) or Paliperidone (which has highest affinity for the 5-HT2A and 5-HT7 serotonin receptors, but can also bind to *α*-adrenergic receptors) (Chue and Chue, 2012; Ebdrup et al., 2011; Jones et al., 2020), did not inhibit tumourigenesis alone or in combination with Trametinib (Fig S5A-D). In support of this, another specific serotonin receptor inhibitor, Ketanserin, targeting the 5-HT2 family (Creed-Carson et al., 2011; Hedner and Persson, 1988), was tested in the primary screen and was not identified as being able to synergise with Trametinib in reducing tumour burden (File S1; data not shown). Furthermore, multiple other serotonin receptor antagonists targeting 5-HT1, 5-HT2, 5-HT3, 5-HT4, 5-HT5, 5-HT6, 5-HT7 families or multiple serotonin receptor types (e.g. Methiothepin mesylate, 3-Tropanylindole-3-carboxylate methiodide, 1-(1-Naphthyl)piperazine hydrochloride, (S)-Propranolol hydrochloride, S(-)-UH-301 hydrochloride, SB 200646 hydrochloride, Cyclobenzaprine hydrochloride, 3-Tropanyl-indole-3-carboxylate hydrochloride, SB 206553 hydrochloride, Metoclopramide hydrochloride, SDZ-205,557 hydrochloride, Granisetron hydrochloride, Ro 04-6790 dihydrochloride, SB 269970 hydrochloride, LY-310,762 hydrochloride, 5-Carboxamidotryptamine maleate, Methiothepin maleate) were tested in the primary screen but did not reduce tumour burden (File S1; data not shown). Thus, these results indicate that the key target for Ritanserin is Dgk*α*, rather than serotonin receptors, in its synergistic effect with Trametinib in reducing Ras-driven polarity-impaired tumourigenesis.

To confirm these findings genetically, we utilized another *Drosophila* model of Ras-driven, polarity-impaired tumourigenesis, that of inducible knockdown of the *dlg1* polarity gene combined with expression of *Ras^V12^* within the whole eye-antennal tissue (Willecke et al., 2011). Using this model, we could induce tumours by outcrossing the stock to a neutral control (*UAS-luciferase*) or to *UAS-RNAi* lines targeting *Dgk* or serotonin receptor genes, and by comparison assess the impact on tumour burden. *Drosophila* has 5 serotonin receptor orthologs, 5-HT1A, 5-HT1B, 5-HT2A, 5-HT2B, and 5-HT7 (Huser et al., 2017). Although *5-HT1B*, *5HT2A*, and *5HT2B* mRNA expression are not detectable in the 3^rd^ instar larval stages (modENCODE RNAseq), we tested the effect of each gene individually using *UAS-RNAi* lines (which were confirmed as effective in knocking down the expression of their corresponding genes in *wild-type* adult/pupal tissue, Fig S6) in the *dlg^RNAi^ /Ras^V12^* eye-antennal epithelial tumours and in *wild-type* eye epithelia (Fig 5). Whilst knockdown of 5-HT2A had a small effect in reducing the size of otherwise *wild-type* eye discs (Fig 5G, quantified in 5S), the knockdown of these genes did not reduce tumour size, and, in fact, 5-HT7 slightly increased tumour size (Fig 5D,F,H,J,L, quantified in 5S). Although the knockdown efficiency of 5-HT2A and 5-HT7 using the RNAi lines was only at ∼40-50% when tested in *wild-type* tissues (Fig S6), the fact that they have these phenotypic effects on tissue size suggests that their knockdown should have been effective enough to reduce tumour burden if they were required for polarity-impaired Ras-driven, tumourigenesis. Thus, these results indicate that, individually, the serotonin receptor genes are not required for polarity-impaired, Ras-driven tumourigenesis, although we cannot formally rule out that they might function redundantly to facilitate tumourigenesis.

**Figure 5.**
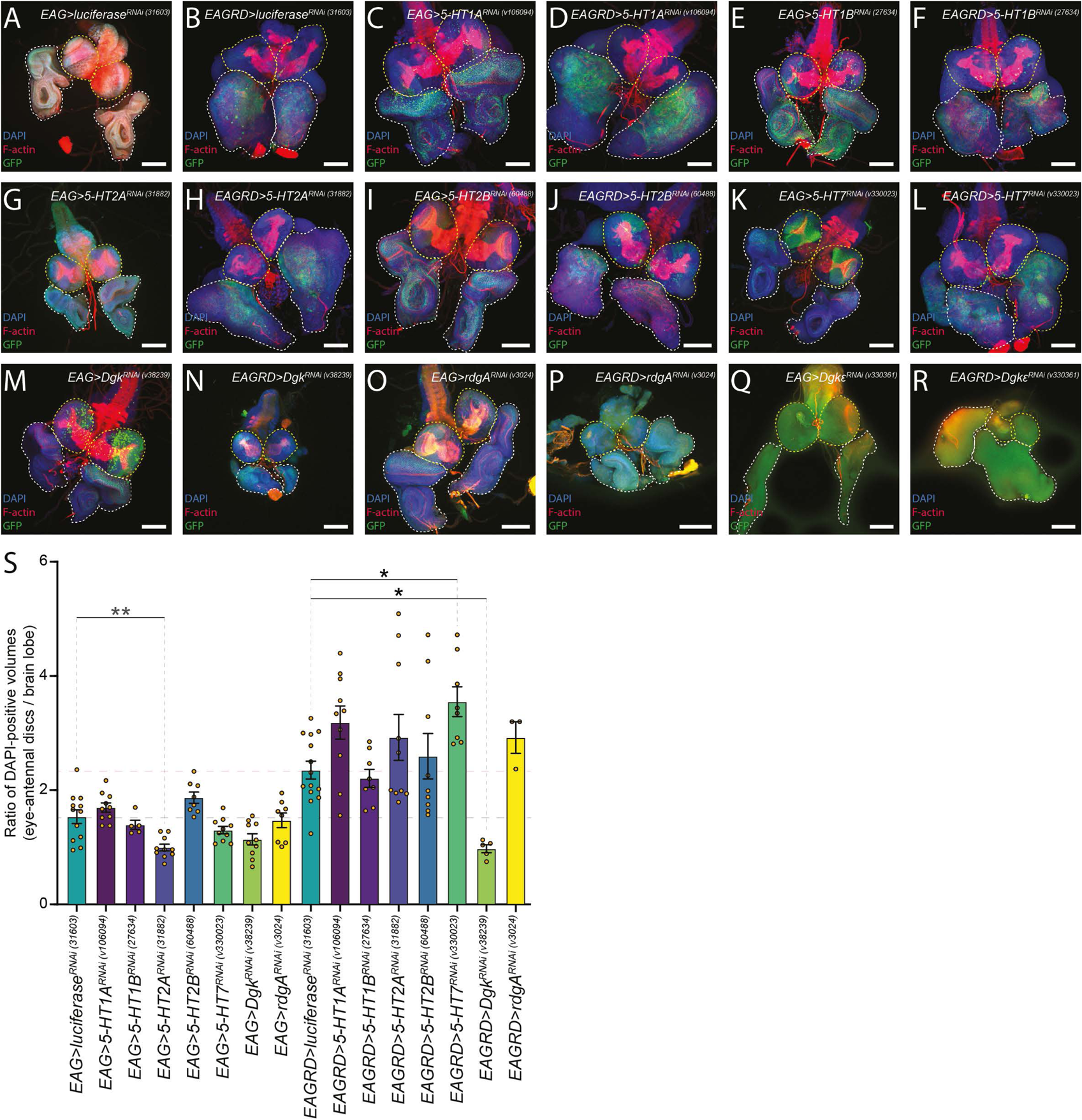
Genetic analyses of *dlg1^RNAi^*/*Ras^V12^* tumours reveals *Dgk* knockdown, but not *Dgkε, rdgA* or serotonin receptor gene expression knockdown, reduces tumour burden. (A-R) Brains from L3 animals with their attached *wild-type* (EAG>, A,C,E,G,I,K,M,O,Q) or *dlg1^RNAi^*/*Ras^V12^*-expressing (EAGRD>, B,D,F,H,J,L,N,P,R) eye-antennal discs, also expressing GFP (green) and RNAi against 5HT (C-L) or DGK family genes, stained with DAPI (blue, except for Q,R) and for F-actin (red). Where possible, the brain lobes are outlined in yellow, and the eye-antennal discs are outlined in white. Scale bars represent 100µm. (S) Quantification of the volume ratio between the eye-antennal discs and their respective brain lobes from samples in A-R. Little change is observed comparing the ratios of EAG> samples. However, in EAGRD> samples, *Dgk* knockdown significantly reduces the eye-antennal disc/brain lobe ratio, suggesting *Dgk* is necessary for tumour growth. Statistical test used was a one-way ANOVA with Tukey’s multiple comparisons, with non-significant comparisons not visualised. Error bars represent S.E.M. * = p<0.05, ** = p<0.01.

There are 3 *Dgk* orthologs in *Drosophila*, *Dgk (*with highest homology to mammalian *DGKα,β,γ)*, *Dgkε* (ortholog of mammalian *DGKε*) and *rdgA* (with highest homology to mammalian *DGKζ,ι*) (Mérida et al., 2008), which are all expressed in 3^rd^ instar larval tissue (modENCODE RNAseq). We tested the effect of knockdown of these genes on *wild-type* eye disc size and *dlg-RNAi Ras^V12^* tumour burden (Fig 5M-R, quantified in 5S), using RNAi lines that were shown to be highly effective at reducing mRNA levels (Fig S6). Of these genes, only the knockdown of *Dgk* expression was able to significantly reduce tumour burden (Fig 5N, quantified in 5S), but also had no effect on *wild-type* eye disc size (Fig 5M, quantified in 5S). Taken together with the results from the drug testing, these data provide evidence that *Dgk*, but not *Dgkε*, *rdgA*, or the serotonin receptor genes, is required for Ras-driven polarity-impaired tumourigenesis.

### Chemical inhibition of DGK reduces survival of *SCRIB* knockdown *H-RAS^G12V^* human mammary epithelial cells

To determine whether our results in *Drosophila* were applicable in human cells, we used the normal mammary epithelial cell line, MCF10A, stably transformed with human *H-RAS^G12V^* and *SCRIB^RNAi^* (Dow et al., 2008) (Fig S7A), and tested whether Ritanserin, R-59-022, or CU-3 were able to synergise with Trametinib in inducing cell death. We first conducted dose response analyses with these drugs to determine their effect on cell survival in MCF10A *SCRIB^RNAi^/H-RAS^G12V^* cells and control MCF10A cells, using the CellTiter-Glo assay, and calculated the IC_50_ for each drug (Fig 6A-D). The MCF10A *SCRIB^RNAi^/H-RAS^G12V^* cells responded similarly to the control cells to each drug alone, except for R-59-022, where the IC^50^ was ∼3.5x higher for the *SCRIB^RNAi^/H-RAS^G12V^* cells (Fig 6A-D). We then treated MCF10A *SCRIB^RNAi^/H-RAS^G12V^* cells and control MCF10A cells with different dose combinations of Trametinib and Ritanserin, R-59-022, or CU-3 approaching the IC_50_ for each drug, and measured cell survival (Fig 6E-J, Supplementary File 3). To determine synergy we used the BLISS independence analysis (Goldoni and Tagliaferri, 2011), where a score >0 indicates synergy. For Ritanserin and Trametinib, while we found synergy between the drugs in both control cells (Fig 6E) and *SCRIB^RNAi^/H-RAS^G12V^* cells (Fig 6F), *SCRIB^RNAi^/H-RAS^G12V^* cells showed higher BLISS scores than control cells. The highest BLISS scores (∼15) in *SCRIB^RNAi^/H-RAS^G12V^* cells occurred at low doses of Trametinib and half IC_50_ doses of Ritanserin (Fig 6F). Examining the effect of Trametinib and Ritanserin on MCF10A *H-RAS^G12V^* and MCF10A *SCRIB^RNAi^* cells revealed that the dose response to the drugs was similar between these cells (Fig S7B, C). Overall, these results show that this drug combination could be used effectively to kill both activated RAS and/or polarity-impaired cells. The combination of R-59-022 and Trametinib (Fig 6G, H) showed overall weak synergism in *SCRIB^RNAi^/H-RAS^G12V^* cells (BLISS scores of ∼10) and in control cells (BLISS scores <10). The DGK*α*-specific inhibitor, CU-3, showed strong synergy with Trametinib in both control (Fig 6I) and *SCRIB^RNAi^/H-RAS^G12V^* cells (Fig 6J), with the highest synergy scores (>25) obtained at approximately half IC_50_ doses of each drug. Altogether, these studies show that the results obtained on the synergy between Trametinib and Ritanserin, R59-022 or CU-3 in inhibiting Ras-driven polarity-impaired tumourigenesis in *Drosophila* are translatable to human cells that have oncogenic RAS and polarity-impairment.

**Figure 6.**
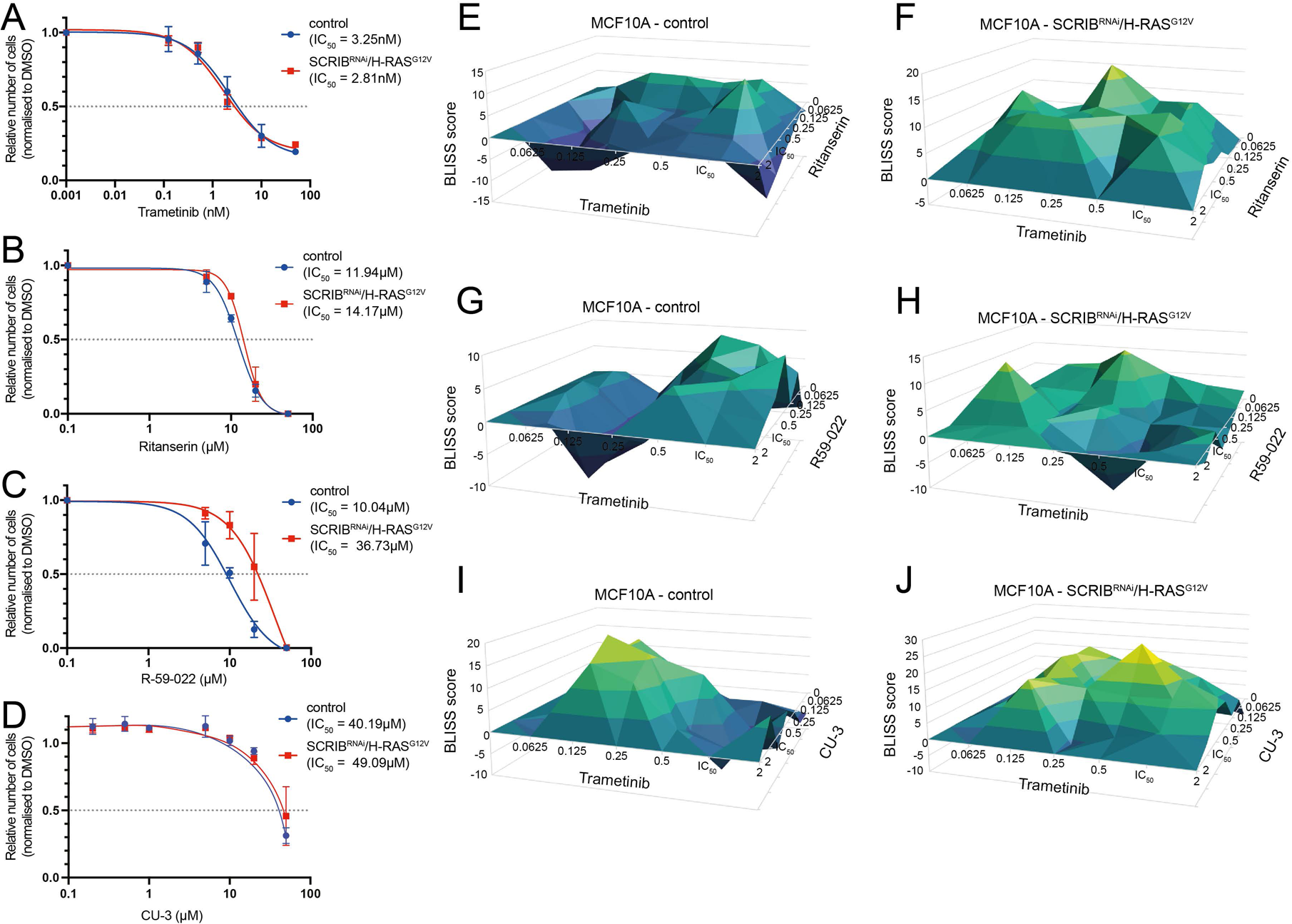
Trametinib synergises with Ritanserin, R-59-022, and CU-3 in cultured MCF10A cells expressing *shRNA-SCRIB* and human *H-RAS^G12V^*. (A-D) Dose response curves were generated for Trametinib (A), Ritanserin (B), R-59-022 (C), and CU-3 (D) treatments in MCF10A cells (both control cells and those with knockdown of *SCRIB* and expression of *H-RAS^G12V^*). Live cell proportions were determined via CellTiter-Glo assay, and IC_50_ values were calculated from 2-3 independent experiments. (E-J) BLISS synergy scores for control (E,G,I) and *shRNA-SCRIB* and human *H-RAS^G12V^* (F,H,J) MCF10A cells treated with Trametinib in combination with Ritanserin (E,F), R-59-022 (G,H), or CU-3 (I,J) at the indicated doses, which are approximate magnitudes of the IC_50_ for each drug. The proportion of live cells was determined by CellTiter-Glo assay, across 3 replicate experiments. A BLISS score > 0 indicates synergy.

### Signalling pathways affected by the Trametinib and CU-3 drug combination

As CU-3 specifically targets DGK*α* (Liu et al., 2016; Yamaki et al., 2019), we focused on the signalling pathways that are known to be regulated by DGK*α*. DGK*α* is one of a family of 10 DGKs that function as lipid kinases phosphorylating diacyl glycerol (DAG) to generate phosphatidic acid (PA), which is an important secondary messenger lipid involved in the regulation of the RAS superfamily (RAS, RAC, ARF) and mTOR (Mechanistic Target of Rapamycin) signalling pathways to control cell growth and proliferation, membrane trafficking and cytoskeletal reorganisation (Andresen et al., 2002; Foster et al., 2014; Zhang and Du, 2009). To determine whether the treatment of *SCRIB^RNAi^/H-RAS^G12V^* cells or control cells with CU-3 + Trametinib inhibited the RAS and mTOR pathways, we treated cells with the drug combination or the vehicle control for 24hr or 48hr and analysed cell lysates for the levels of the mTOR target, pS6K, versus total S6K protein, and the levels of the MEK target, pERK, versus total ERK (Fig 7). For this experiment, we chose the concentration of CU-3 (10 uM) and Trametinib (1.515 nM) that showed the most effective synergistic effects in the BLISS assay (Fig 6J). As expected, we found that pERK levels were upregulated in *SCRIB^RNAi^/H-RAS^G12V^* cells compared with control cells and upon combination drug treatment were reduced by ∼4-fold at 24hr and 48hr, whereas in control cells the effect was much less (∼2-fold reduction). Similarly, pS6 levels were elevated in *SCRIB^RNAi^/H-RAS^G12V^* cells relative to control cells, and the combination drug treatment dramatically reduced pS6 levels (by ∼15-fold at 24 hr and by ∼3-fold at 48 hr), but only showed a 2-fold decrease in control cells at 24hr. Thus, as expected the CU-3 + Trametinib treatment leads to an inhibition of RAS-MEK-ERK and mTOR-S6K signalling. Since mTOR can also be positively regulated by PI3K signalling and the PI3K pathway can be upregulated by RAS signalling (Carnero, 2010; Memmott and Dennis, 2009), we also analysed the levels of a target of the PI3K target, pAKT, relative to total AKT protein (Fig 7). Interestingly, pAKT levels were not greatly affected in *SCRIB^RNAi^/H-RAS^G12V^* cells or in control cells at 24hrs or 48hr, and the combination drug treatment did not affect pAKT levels at 24hr, but at 48hr there was an increase pAKT levels ∼2-fold in the *SCRIB^RNAi^/H-RAS^G12V^* cells and 3-fold in the control cells. This increase in pAKT in the *SCRIB^RNAi^/H-RAS^G12V^* cells and control cells in response to the combination drug treatment might be due to a compensatory response to the inhibition of mTOR signalling due to the disruption of a negative feedback loop between mTOR and PI3K that has been previously documented (Harrington et al., 2005; Manning et al., 2005). Altogether, our signalling pathway analyses reveal that combined inhibition of DGK*α* and Ras signalling in *SCRIB^RNAi^/H-RAS^G12V^* cells results in the reduction of pERK and pS6K levels at an early stage (24hr) after drug exposure but does not affect pAKT at this timepoint, indicating that the drug combination targets the Ras and mTOR signalling pathways but not the PI3K pathway.

**Figure 7.**
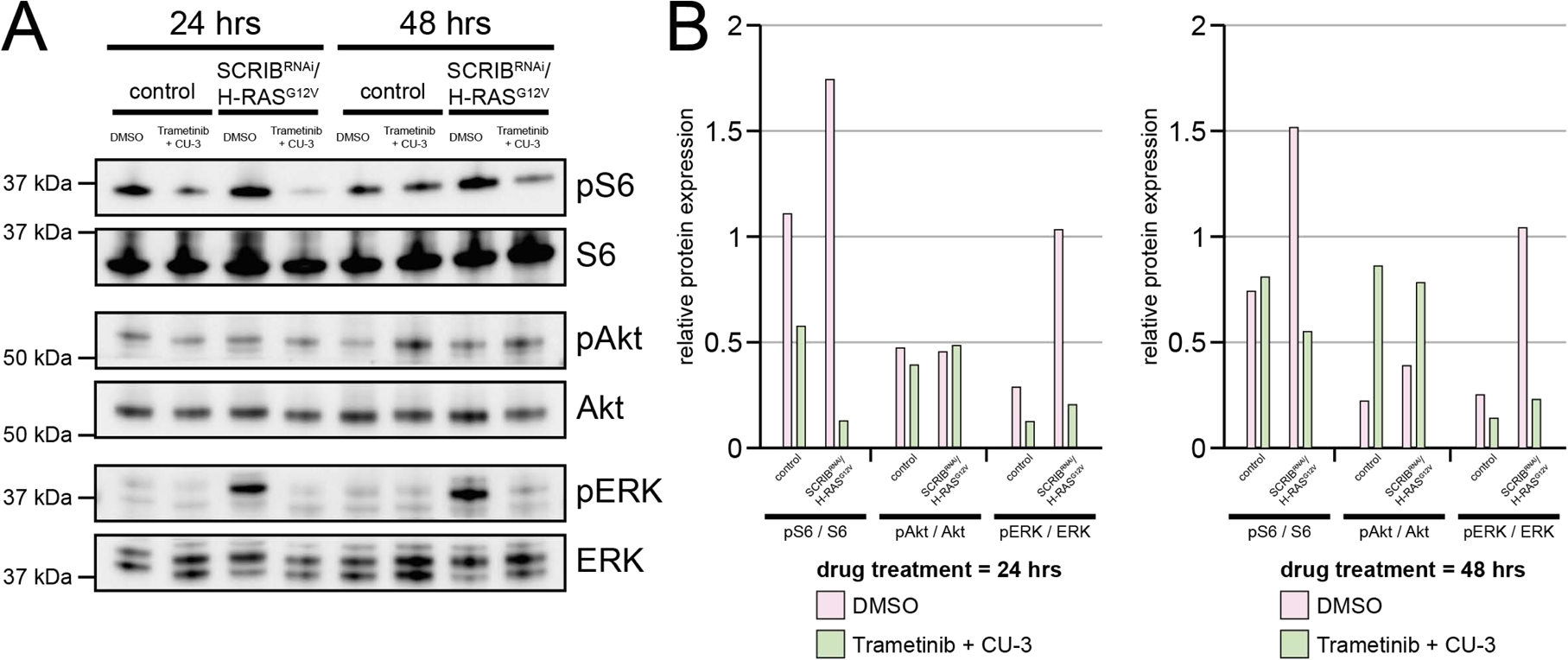
The specific DGKα inhibitor CU-3 synergizes with Trametinib by inhibiting the mTOR and Ras signalling pathways. (A) WB images of MCF10A control or *SCRIB^RNAi^/H-RAS^G12V^* cells, treated for either 24 or 48 hrs with either DMSO or Trametinib and CU-3 in combination. (B) Quantifications of WBs in A, measuring phosphorylated protein levels relative to the unphosphorylated protein. Relative to the respective DMSO control, pS6 and pERK levels are reduced in samples treated with Trametinib and CU-3, while pAkt levels are increased (likely due to feedback from mTOR inhibition).

## Discussion

Combination drug therapy is becoming increasingly favoured as an anti-cancer therapy, as it can allow lower doses of each drug to be used, decreasing unwanted side effects and reducing the chance drug resistance will occur (Al-Jundi et al., 2020; De Leo et al., 2020; Ferrari et al., 2020; Roskoski, 2017). However, identifying novel drug combinations that are effective against specific cancers can be a difficult prospect. In this regard, the use of the *Drosophila* model has become a highly effective *in vivo* tool in discovering bioavailable efficacious drug combinations that are translatable to human cells, and in some cases also clinically effective as anti-cancer therapies (Bangi et al., 2019; Bangi et al., 2016; Levine and Cagan, 2016; Stickel et al., 2015). Using a *Drosophila* model of Ras-driven, polarity-impaired cancer, we undertook a boutique screen of chemical libraries and identified two compounds that showed synergy with subtherapeutic doses of the MEK inhibitor, Trametinib, identifying the Polo-like kinase inhibitor, Volasertib, and the 5-HT2A/2C serotonin receptor and DGK*α* inhibitor, Ritanserin. Polo-like kinase inhibitors have been extensively utilised in cancer therapy (Gjertsen and Schöffski, 2014; Shakeel et al., 2021; Zhang et al., 2021), and have also been explored as a potential combination therapy with MEK inhibitors for the treatment of *N-RAS*-driven melanoma (Posch et al., 2015). Thus, we focused on characterizing the synergistic interaction between Trametinib and Ritanserin, revealing by pharmacogenetic analysis in *Drosophila* that the key target of Ritanserin is Dgk (ortholog of mammalian DGK*α/β/γ*) rather than other *Drosophila* Dgk paralogs or serotonin receptors. Furthermore, we show that our findings in *Drosophila* are translatable to MCF10A human mammary epithelial cells harbouring oncogenic *H-RAS* and knockdown of the cell polarity gene, *SCRIB*. We demonstrated that both Ritanserin and another serotonin receptor and DGK*α* inhibitor, R-59-022, synergised with low-doses of Trametinib in *SCRIB^RNAi^/H-RAS^G12V^* cells, as well as in *SCRIB^RNAi^* cells and *H-RAS^G12V^* cells at doses where normal cells were not greatly affected, thus suggesting that these drug combinations could be used to specifically target Ras-driven and/or polarity-impaired cancer cells. Using a DGK*α* specific inhibitor, CU-3, we also demonstrated synergy with Trametinib in MCF10A cells, but in this case drug dose combinations affected the *SCRIB^RNAi^/H-RAS^G12V^* cells similarly to normal cells, suggesting that CU-3 would not be clinically useful in specifically targeting Ras-driven cancer cells. However, another compound that targets DGK*α/β/γ* isoforms (Compound A) has been identified as highly effective in inducing apoptosis of a variety of cancer cells *in vitro* (Yamaki et al., 2019), and it would be interesting to determine whether this compound, as well as other newly identified DGK-specific inhibitors (Velnati et al., 2020; Velnati et al., 2019), may have greater efficacy with Trametinib in specifically targeting Ras-driven cancer cells relative to normal cells. Of the DGK*α* (and 5-HT2A/2C serotonin receptor) inhibitors we analysed, only Ritanserin has been tested in human clinicals trials, but in this case for schizophrenia, cocaine and alcohol dependence, and migraines (Cornish et al., 2001; Johnson et al., 1996; Nappi et al., 1990; Wiesel et al., 1994), though it has not been marketed for clinical use. *In vitro* studies have shown that Ritanserin induces cell death in the mesenchymal cell subtype of glioblastomas (Audia and Bhat, 2018; Olmez et al., 2018). Furthermore, it was shown that Ritanserin can synergize with irradiation and with Imatinib, a tyrosine kinase inhibitor, to decrease cell proliferation of mesenchymal glioblastoma cells (Olmez et al., 2018). Similarly, the DGK*α*-specific inhibitor, Amb639752, promotes restimulated apoptosis of cells *in vitro* and in animal models of X-linked lymphoproliferative disease type 1, which have elevated DGK*α* activity (Velnati et al., 2021a; Velnati et al., 2021b; Velnati et al., 2020). These studies, along with our own findings, suggests that Ritanserin, as well as other DGK*α* inhibitors, would be good candidates to be developed clinically as anti-cancer drugs and to be used in combination with Trametinib or other EGFR-RAS pathway inhibitors, particularly in Ras-driven cancers.

DGK*α* is considered an oncogene, being upregulated in many cancers, and promotes cell proliferation and cell survival (Chen et al., 2019; Fazio et al., 2020; Merida et al., 2017; Sakane et al., 2021). Mechanistically, we showed that inhibition of DGK*α* in human mammary epithelial cells harbouring the *H-RAS* oncogene and knockdown of the cell polarity gene, *SCRIB*, resulted in reduced MEK and mTOR activity. Elevated DGK*α* activity phosphorylates diacyl glycerol (DAG) to form phosphatic acid (PA), promoting signalling pathways regulated by PA, and attenuating those regulated by DAG (Andresen et al., 2002; Foster et al., 2014; Zhang and Du, 2009) (Fig 8). PA upregulates RAS-RAF-MEK-ERK signalling by increasing the binding of cRAF to the endosomal membranes, where it interacts with active RAS (RAS-GTP) and forms a scaffold with MEK and ERK, thereby enabling RAS pathway activation and promoting cell proliferation and survival (Andresen et al., 2002). PA also promotes mTOR activity by enabling the interaction of mTOR with the Raptor adaptor protein to form the mTORC1 complex, and with the Rictor adaptor protein to form the mTORC2 complex (Fang et al., 2001; Toschi et al., 2009) (Fig 8). DGK*α* has also been reported to positively regulate mTOR transcription through a mechanism involving phosphodiesterase 4A1 and cyclic-AMP (Dominguez et al., 2013). mTOR, through its promotion of protein synthesis, plays an important role in stimulating cell growth and proliferation (Saxton and Sabatini, 2017). Thus, by inhibiting DGK*α* activity and decreasing PA, RAS and mTOR signalling will be inhibited, thereby inhibiting cell growth, proliferation and survival. The identification of mTOR signalling as a pathway that was inhibited in *SCRIB^RNAi^/H-RAS^G12V^* cells that were growth inhibited upon treatment with CU-3 + Trametinib is consistent with *Drosophila* genetic studies showing that the mTOR pathway is important for polarity-impaired Ras-driven tumourigenesis (Willecke et al., 2011). In mammalian systems, the interaction of SCRIB with PI3K pathway components, PHLPP1 and PTEN, negatively regulates the PI3K-AKT-mTOR pathway, and upon SCRIB downregulation or mutation pathway activation is observed (Feigin et al., 2014; Li et al., 2011), but whether PI3K-AKT-mTOR pathway activation is required for tumourigenesis upon *SCRIB* downregulation or inactivation in mammalian cells is not known. By contrast, RAS signalling is negatively regulated by *SCRIB* and is required in *SCRIB* knockdown mammalian cells for hyperplastic growth *in vitro* and *in vivo* (Dow et al., 2008; Nagasaka et al., 2010; Nagasaka et al., 2013; Pearson et al., 2011; Stephens et al., 2018).

**Figure 8.**
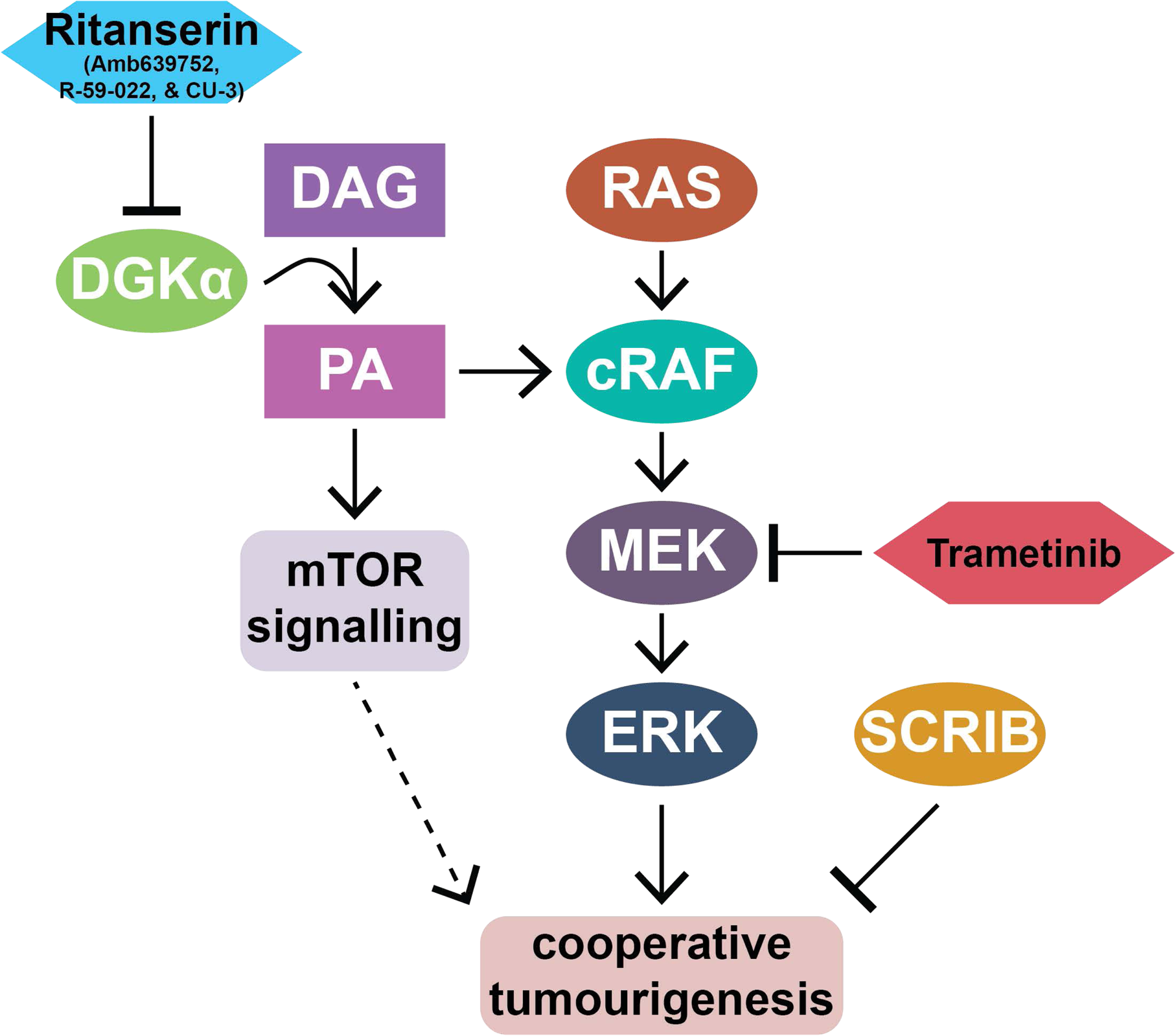
Model of DGK involvement in *scrib^-^/Ras^V12^* tumourigenesis, and the effect of Ritanserin and Trametinib. Activated RAS induces signalling via the cRAF-MEK-ERK cascade, promoting hyperplastic tissue growth. The additional inhibition of SCRIB results in cooperative tumourigenesis. Trametinib, a MEK inhibitor, can modulate this signalling and leads to a reduction in tumour growth. Our data show that Trametinib synergises with Ritanserin to strongly inhibit tumourigenesis. Ritanserin acts by inhibiting the activity of DGK*α*, which catalyses the conversion of diacylgycerol (DAG) into phosphatidic acid (PA). Evidence suggests that PA stimulates cRAF-MEK-ERK signalling, as well as mTOR signalling, which are downregulated in tumourigenic cells treated with Trametinib and CU-3, a DGK*α*-specific inhibitor. Other acronyms: RAS (RAS proto-oncogene, GTPase), cRAF (Raf-1 proto-oncogene, serine/threonine kinase), MEK (mitogen-activated protein kinase kinase 7), ERK (mitogen-activated protein kinase 1), SCRIB (scribble planar cell polarity protein), mTOR (mechanistic target of rapamycin kinase), DGK*α* (diacylglycerol kinase alpha).

Whether other pathways known to be regulated by DGK*α* (Sakane et al., 2021) are important targets in *SCRIB^RNAi^/H-RAS^G12V^* cells will be important to investigate. One such pathway, activated by PA, is atypical protein kinase C zeta (PKC*ζ*) – Nuclear factor kappa B (NF*κ*B) signalling, which promotes cell survival (Kai et al., 2009; Sakane et al., 2021). Atypical protein kinase C (aPKC), is also a cell polarity regulator that plays important roles in establishing and maintaining apical-basal cell polarity of epithelial cells (Tepass, 2012; Vorhagen and Niessen, 2014), and also negatively regulates the Hippo negative tissue growth control signalling pathway in both mammalian cells (Archibald et al., 2015) and *Drosophila* (Doggett et al., 2011; Grzeschik et al., 2010). As *Drosophila* polarity-impaired Ras-driven tumours have unrestrained aPKC activity, an important factor in inducing tumour growth (Leong et al., 2009), which occurs via Hippo pathway inhibition (Doggett et al., 2011), it is possible that by inhibiting DGK*α,* the elevated aPKC activity and Hippo pathway impairment will be rescued. *Drosophila* polarity-impaired Ras-driven tumours also upregulate the JNK stress response pathway to promote tumourigenesis by blocking differentiation and promoting invasion, and in mammalian *SCRIB*-impaired cells the P38 stress response pathway is elevated (Norman et al., 2012). It will therefore be important to determine whether DGK*α* inhibition also alters these signalling pathways.

Another important aspect of DGK activation is that DAG levels will be depleted, which has the potential to affect various signalling pathways (Griner and Kazanietz, 2007; Merida et al., 2008). DAG binds to and activates the C1 domain of the Protein Kinase C family proteins, as well as other non-kinase proteins including GTPase regulators (Colon-Gonzalez and Kazanietz, 2006; Griner and Kazanietz, 2007; Oliva et al., 2005; Wang and Kazanietz, 2006). Protein Kinase C family members function downstream of growth factor receptors to promote signalling (Oliva et al., 2005), and therefore DGK activation and decreased DAG would limit their activation. One of the non-kinase proteins activated by DAG is RAC-GAP, which is involved in the inactivation of the RAC-GTPase (Wang and Kazanietz, 2006), an important regulator of the actin cytoskeleton and signalling from cell-cell adhesion junctions (Bosco et al., 2009; Ratheesh et al., 2013). DGK activation and lower DAG would therefore reduce RAC-GAP activity and increase RAC activity, which might contribute to *SCRIB^RNAi^/H-RAS^G12V^* tumourigenic properties, as *Drosophila* studies have shown that Rac activates JNK signalling (Brumby et al., 2011; Ma et al., 2015), a key factor in promoting *scrib* mutant Ras-driven tumourigenesis (Igaki et al., 2006; Leong et al., 2009; Uhlirova and Bohmann, 2006). DGK*α* inhibition, by increasing DAG and activating Rac-GAP, would be expected to reduce Rac activity and JNK activity and inhibit *scrib* mutant Ras-driven tumourigenic properties. However in mammalian epithelial cells, the effect of RAC on tumourigenesis may have a different role, since SCRIB binds to the RAC regulators, *β*PIX and GIT1, and is involved in the activation of RAC in cMYC-transformed cells to induce JNK-mediated cell death, but in *SCRIB* depleted cells RAC activity is reduced thereby promoting cell survival (Zhan et al., 2008). DGK*α* inhibition, by increasing DAG and activating RAC-GAP, would be expected to reduce RAC activity even further in *SCRIB* depleted cells, but whether this will affect *SCRIB^RNAi^/H-RAS^G12V^* tumourigenesis similarly to the situation in *Drosophila* remains to be determined. The investigation of the involvement of DAG-regulated signalling pathways in *SCRIB^RNAi^/H-RAS^G12V^* tumourigenesis, and the effect of inhibition of DGK*α* on these pathways, will be important in providing a full picture of the involvement of DGK*α* in RAS-driven cancers.

In summary, our analyses have revealed that DGK*α* plays an important role in Ras-driven polarity-impaired tumourigenesis in both the *Drosophila* model and mammalian epithelial cell lines. Our findings raise many questions that remain to be investigated, such as whether DGK*α* activity is elevated in Ras-driven polarity-impaired cells and, if so, how this occurs, and whether increased PA-signalling or decreased DAG signalling provide the critical function of DGK*α* in tumourigenesis. However, since DGK*α* is upregulated and oncogenic in human cancer (Chen et al., 2019; Fazio et al., 2020; Merida et al., 2017; Sakane et al., 2021), the findings from our study suggest that Ras-driven polarity-impaired cancers may be particularly addicted to DGK*α* for tumour survival, and suggests that DGK*α*-inhibitors combined with Ras pathway inhibitors will be a highly effective drug combination for anti-cancer therapy in these cancers. DGK*α* inhibitors have been considered for development as anti-cancer therapy not only for their effect on the cancer but also on the T-cell anti-cancer immune response (Arranz-Nicolas and Merida, 2020; Sakane et al., 2008; Sakane et al., 2016). Our findings suggest that combining Ras pathway inhibitors with DGK*α* inhibitors may provide even greater efficacy against Ras-driven cancers, as well as decrease unwanted side-effects and reduce the development of drug resistance.

## Materials and Methods

### Drosophila melanogaster stocks and husbandry

Fly stocks used in this study are detailed in Table S1. Unless otherwise indicated, animals were maintained, and crosses were undertaken on a standard cornmeal/molasses/yeast medium within temperature-controlled incubators at 25°C.

### Generation of larvae for screening

Approximately 450 *eyFLP, UAS-GFP;; tub-GAL4, FRT82B, tub-GAL80 / TM6B, Tb-RFP* females were crossed to approximately 150 *UAS-Ras85D^V12^, FRT82B, scrib^1^ / TM6B* males. Crosses were undertaken in cages containing apple juice agar plates (“lay plates” – 35g agar (Amresco #J637) and 20g sucrose (SIGMA #S-0389) dissolved in 1L H_2_O by microwaving, add 250mL apple juice (Spring Valley), incubate at 60⁰C for 1-2 hrs, add 25mL Tegosept solution (100g methylhydroxybenzoate in 1L abs. ethanol), then set in 10cm Petri dishes). On the lay plates, larvae/flies were fed with yeast paste (∼100g compressed yeast (Lesaffre) in H_2_O to thickened consistency). A laying period of 7 – 14 hours was used. Early third-instar larvae (as indicated by size) with the genotype *eyFLP, UAS-GFP; UAS-Ras85D^V12^ / + ; FRT82B, scrib^1^ / tub-GAL4, FRT82B, tub-GAL80* were collected at 48-60 hours after egg laying by rinsing the lay plate with tap water before tipping the contents through a small sieve. Larvae were then selected for compound screening plates based on the presence/absence of physical markers (observed GFP expression in head region, and lack of TM6B-RFP markers), using a SteREO Discovery.V8 microscope (Zeiss). Method adapted from Willoughby et al. (2013) (Willoughby et al., 2013).

### Compound screening

Approximately 2mg of instant *Drosophila* media (Southern Biological #CM4) was added to each well of a deep-well 96-well plate (NUNC #260252). This media was reconstituted with 240µL yeast solution (14g dried yeast (Tandaco) dissolved in 250mL H_2_O by microwaving for 10 mins to inactivate yeast, and stored at 4⁰C), which contained the compound(s) of interest (File S1), or a DMSO control at 0.5% v/v. Larvae (generated as above) were then added to each well of the 96-well plate at a density of ∼7 per well. The plate was sealed with wire mesh and perspex (containing 96 holes for airflow), and incubated at 25⁰C for 5 days (∼120 hrs). An unsealed container of water was incubated next to the plate to maintain humidity and prevent desiccation. Sucrose solution (30% w/v sucrose in H_2_O) was then added to each well and the larvae dislodged from the food via agitation. Additional sucrose solution was added to each well until a convex meniscus formed, allowing larvae to float to the top of the well and into the focal plane. The plate was then imaged 2 wells at a time using the SteREO Discovery.V8 microscope with Zen 2012 software (Zeiss). The resulting images were “stitched” together using Photoshop (various editions, Adobe). The stitched image was then thresholded using Fiji (Schindelin et al., 2012), with the pixel intensity threshold determined as the value that first eliminated any background signal. This resulted in a binarized image where white pixels represented GFP-positive tumourigenic tissue. A white pixel count was then performed for each well using the Fiji ‘Analyse Particles’ function (Schindelin et al., 2012) To maximise consistency results, an ImageJ Macro (Mutterer and Rasband, 2012) was written, which automatically performed white pixel counts automatically on areas corresponding to the plate wells, using a nested FOR loop (Supplementary File 4). Areas of white pixels below 2 pixels were excluded from the analyses as noise. The data was then exported to Excel (Microsoft), and the pixel count for each well was divided by the number of larvae in the respective well, producing the mean GFP-positive pixel area per organism for each well as a representation of tumour size.

### Compounds

Compounds utilised in this study were selected on the basis of their identification as potential hits in a primary compound screen (unpublished data). The primary compound screen was performed as described above. Each compound derives from one of four boutique screening libraries (“epigenetic library”, “kinase library”, “targeted agents library”, and “FDA-approved known drug library”) obtained from Hélène Jousset Sabroux and Kym Lowes (WEHI, Australia). All compounds were dissolved in DMSO, then 2mL yeast solution, to obtain the desired concentrations (see File S1, Table 1 and relevant figures), and such that the final concentration of DMSO was not greater than 0.5% v/v. The following additional compounds were obtained from the following sources: R-59-022 (CAYMAN CHEMICAL COMPANY #16772), Amb635792 (AMBINTER #Amb635792) and CU-3 (Med Chem Express #HY-121638A).

### Statistical analyses

For each compound treatment, for each well, the mean GFP-positive pixel area per organism was divided by the mean GFP-positive pixel area per organism of all DMSO-treated wells, obtaining the “mean GFP pixel intensity per organism per well normalised to DMSO”. These values were analysed statistically using Prism (GraphPad), with the particular tests employed for each analysis detailed in the respective figure legends.

### GFP-positive area to larvae pixel area ratio analysis

Binarized plates images and their respective unaltered plate images were imported into Photoshop 2020 (Adobe). For 2 larvae per well, larval size (area in pixels) was determined using the “Lasso” tool. The GFP-positive tumour size (area in pixels) for the respective tumour(s) of each larvae measured were determined using the “Magic Wand” tool. The GFP-positive pixel area to larvae pixel area ratio was then calculated for each animal by dividing the size of the tumour by the size of the respective larva.

### RNA extraction, cDNA synthesis, and qRT-PCR

RNAi lines were crossed to *hsFLP ; Actin>>GAL4, UAS-GFP* and raised at 25⁰C. Whole adults or pupae (as some crosses were lethal at the pupal stage) were homogenised (n=∼10 animals per genotype) in 1× phosphate buffered saline (PBS). RNA extraction and cDNA synthesis was performed as previously described (La Marca et al., 2019). qRT-PCR was performed using a Power SYBR Green PCR Master Mix (Applied Biosystems #4367659) and a QuantStudio 12K Flex Real-Time PCR System (Applied Biosystems). The data were normalised to expression of the housekeeping genes *Gapdh2* and *Rpl32*. The primer sequences used are listed in Table S2, and were obtained from Integrated DNA Technologies.

### Tissue imaging

RNAi lines were crossed to both eyFLP ;; Act>>GAL4, UAS-GFP / TM6B (EAG) and eyFLP ; UAS-Ras85D^V12^, UAS-dlg1 RNAi / CyO tub-GAL80 ; Act>>GAL4, UAS-GFP / TM6B (EAGRD), and incubated at 25⁰C for ∼7 days. UAS-luciferase RNAi (31603) was used as a non-targeting control. 3^rd^ instar larvae were dissected in 1× PBS (AMRESCO #E703), fixed with 4% paraformaldehyde (Alfa Aesar #43368) in 1× PBS with 0.1% Triton X-100 (PBST). Samples were incubated in DAPI (stock prepared at 1μg/mL, used at 1:1000, Sigma-Aldrich #D9542) and phalloidin-tetramethylrhodamine isothiocyanate-Rhodamine solution (used at 0.3µM, Sigma-Aldrich #P1951) to mark DNA and F-actin. Samples were imaged via confocal microscopy using an LSM 780 (Zeiss), and the images processed using Zen 2012 (Zeiss) and Photoshop (Adobe). Imaris (Bitplane) was used to measure the volumes of the eye-antennal discs and brain lobes, which were identified using DAPI-positive tissue.

### MCF10A cell culture

Stably-transformed MCF10A cells were used for cell culture experiments, with the following genotypes: *wild-type* (control), *MSCV-Scramble*, *MSCV-shSCRIB7*, *H-Ras^G12V^-Scramble*, and *H-Ras^G12V^-shSCRIB7* (Dow et al., 2008). MCF10A cells were maintained at 37⁰C and 5% CO2 in DMEM:F12 with donor horse serum (20ng/mL), EGF (100ng/mL), and cholera toxin (100ng/mL).

### Western blotting

For the western blotting experiments, MCF10A cells were plated at 140000 cells/well of a 6-well plate (COSTAR #3506) and incubated at 37⁰C with 5% CO_2_. The next day, media was removed and DMSO or Trametinib (1.1515µM) and CU-3 (10µM) were applied in 1.5mL media. Cells were treated in drugs for 24 hrs or 48 hrs, and then collected by washing with tissue culture grade PBS (TC-PBS), dissociation with 1× trypsin (Lonza #BE02-007E), washing again with TC-PBS, and the pellet collected and snap-frozen, before storage at −20⁰C.

Protein was isolated by incubation for 30 mins on ice in RIPA buffer (150mM NaCl, 0.1% w/v SDS, 1% Triton X-100, 0.5% sodium deoxycholate, 50mM TRIS buffer pH 8.0), with cOmplete Protease Inhibitor Cocktail (Roche #11836153001) as a protease inhibitor and PhosSTOP (Roche #04906845001) as a phosphatase inhibitor. Cell lysate was then centrifuged at 13000rpm for 10 mins at 4⁰C, and the supernatant stored at −20⁰C as needed. For all samples, to determine protein concentrations, a Pierce BCA Protein Assay kit (ThermoFisher #23225) was used. 4× Laemmli buffer (0.25M TRIS-HCl buffer, 40% glycerol, 8% SDS, 0.1% w/v bromophenol blue) with 1:10 β-mercaptoethanol (BDH #441433A) was added to protein lysate and samples boiled for 5 mins. Equal quantities of protein (10-20µg) were loaded on a 4-12% NuPAGE Bis-Tris gel (ThermoFisher #NP0335BOX) and run at 120-150V. Precision Plus Protein Kaleidoscope Prestained Protein Standard (Bio-Rad #1610375) was used as a molecular weight marker. Gels were transferred to an iBlot Transfer Stack nitrocellulose membrane (Life Technologies #IB23001) using an iBlot2 Transfer device as per the manufacturer’s protocols. Membranes were blocked in 5% bovine serum albumin (BSA) (Sigma-Aldrich #A7906) in 1× TBST (1× TRIS buffered saline + 0.1% Tween-20 (Sigma-Aldrich P1379)) for 1 hr at room temperature with gentle agitation. Membranes were then incubated in primary antibodies (Table S3) diluted in 5% BSA in 1× TBST overnight at 4°C with rolling agitation. Membranes were then washed 4 times for 10 mins each in 1× TBST, incubated in the appropriate secondary antibody diluted in 5% BSA in 1× TBST for 1 hr at room temperature, then washed again 4 times for 10 mins each in 1× TBST. Secondary antibodies used were Goat Anti-Mouse Ig, Human ads-HRP (Southern Biotech #1010-05) and Goat Anti-Rabbit Ig, Human ads-HRP (Southern Biotech #4010-05). Immobilon Forte Western HRP Substrate (Merck Millipore #WBLUF0500) was used to resolve the staining, before the membranes were imaged on a ChemiDoc MP Imaging System (Bio-Rad). As needed, membranes were incubated in HRP inactivation solution (0.2% NaN3 in 1× PBS for 30 mins with agitation) or mild stripping solution (glycine 1.5% w/v, SDS 0.1% w/v, Tween-20 1% v/v, pH2.2 in H_2_O) to re-probe the membrane. Western blot images were quantified using Photoshop (Adobe).

### IC_50_ and BLISS assays

MCF10A cells were plated at 1000 cells/well of a 96-well flat-bottom white plate (Greiner CELLSTAR #655083) and incubated at 37⁰C with 5% CO_2_. Wells at the border of the plate were filled with H_2_O to prevent evaporation. The next day, cells were treated with Ritanserin, Trametinib, R-59-022, CU-3, or drug combinations at the concentrations indicated in the relevant figures. Cells were then incubated for a further 72 hrs. Media was then removed from wells and cells were washed with TC-PBS. 40µL TC-PBS was then added to each well, followed by 40µL CellTiter-Glo 2.0 Viability Assay Reagent (Promega #G9241). Plates were incubated on an orbital shaker, in darkness, for 5 mins at room temperature before luminescence was read on a CLARIOstar Plus (BMG Labtech). Luminescence readings were normalised to wells containing DMSO-treated cells. For IC_50_ assays, cells were plated in duplicate, and 2-3 independent experiments were performed, and each IC_50_ was calculated in Prism (GraphPad) using nonlinear regression (curve fit) analysis. For BLISS assays, 2-4 independent experiments were performed and the average reading across experiments used to build the final synergy scores. BLISS synergy scores were calculated for each well using standard methods (Bliss, 1939) and 3D plots generated in Microsoft Excel.

## Acknowledgements

We thank Lee Willoughby for technical advice and all laboratory members for discussions of this projects. We thank Hélène Jousset Sabroux (WEHI) and Kym Lowes (head of WEHI screening laboratory) for advice on the compound libraries. We are grateful to the Vienna *Drosophila* Research Center and Bloomington Stock Center for *Drosophila* stocks and to Flybase for its wealth of information.

## Author contribution

HER, POH designed the project; JELM, RWE, STD, PB conducted experiments; JELM, RWE, STD, PB, HER, POH, GLK analysed data; HER, JELM, RWE wrote the manuscript; JELM, RWE prepared figures; POH, GLK, STD provided editorial advice.

## Supplementary information

**Supp File 1 – List of all drugs screened for synergy with Trametinib in reducing *scrib^-^ Ras^V12^* tumour burden.**

Drugs highlighted in yellow are ones identified as reducing tumour burden in the primary screen that were further explored in this study.

**Supp File 2 – Hits from the primary screen that did not significantly reduce *scrib^-^ Ras^V12^* tumour burden upon retesting.**

Binarized images of the screening plates and graphs of pixel intensity are shown.

**Supp File 3 – Drug combination cell viability analysis.**

**Supp File 4 – Code used to perform automated analyses of binarized images of 96 well plates containing larvae treated with drugs of interest.**

**Table S1 - Fly stocks used in this study.**

**Table S2 - Primers used in qRT-PCR analysis.**

**Table S3 - Primary antibodies used for Western Blots.**

### Supplementary Figure Legends

**Supplementary Figure 1. Methotrexate and Pralatrexate reduce *scrib^-^/Ras^V12^* tumour burden independently of Trametinib.**

(A) Binarized image of a drug test plate where larvae were treated with Trametinib in combination with the folate inhibitor, Methotrexate. (B) Quantifications of GFP pixel intensity revealed that Methotrexate reduces tumour burden, but does not synergise with Trametinib, regardless of the concentration tested. Data from 1 experiment with 16 replicate wells for each treatment. (C) Binarized image of a drug test plate where larvae were treated with Trametinib in combination with another folate inhibitor, Pralatrexate. (D) Quantifications of GFP pixel intensity revealed that Pralatrexate also reduces tumour burden, but does not synergise with Trametinib, regardless of the concentration tested. Data from 1 experiment with 8 replicate wells for each treatment. Statistical tests used were one-way ANOVAs with Tukey’s multiple comparisons. Error bars represent S.E.M. ** = p<0.01, *** = p<0.001, **** = p<0.0001.

**Supplementary Figure 2. Temsirolimus and GSK2126458 reduce *scrib^-^/Ras^V12^* tumour burden independently of Trametinib.**

(A) Binarized image of a drug test plate where larvae were treated with Trametinib in combination with the mTOR inhibitor, Temsirolimus. (B) Quantifications of GFP pixel intensity revealed that Temsirolimus reduces tumour burden, but does not synergise with Trametinib, regardless of the concentration tested. Data from 1 experiment with 16 replicate wells for each treatment. (C) Binarized image of a drug test plate where larvae were treated with Trametinib in combination with the PI3K inhibitor, GSK2126458. (D) Quantifications of GFP pixel intensity revealed that GSK2126458 also reduces tumour burden, but does not synergise with Trametinib. Data from 1 experiment with 8 replicate wells for each treatment. Statistical tests used were one-way ANOVAs with Tukey’s multiple comparisons. Error bars represent S.E.M. ** = p<0.01, *** = p<0.001, **** = p<0.0001.

**Supplementary Figure 3. Reduced Trametinib concentration precludes the synergistic effect.**

(A) Binarized image of a drug test plate where larvae were treated with Trametinib in combination with the Ritanserin, using reduced concentrations of Trametinib. (B) Quantifications of GFP pixel intensity revealed that lower Trametinib concentrations remove the synergistic effects observed together with Ritanserin. Data from 1 experiment with 8 replicate wells for each treatment. Statistical tests used were one-way ANOVAs with Tukey’s multiple comparisons. Error bars represent S.E.M. * = p<0.05, ** = p<0.01.

**Supplementary Figure 4. Other DGK and 5-HT serotonin receptor inhibitory drugs replicate the *scrib^-^/Ras^V12^* tumour reduction obtained using Ritanserin.**

(A) Binarized image of a drug test plate where larvae were treated with Trametinib in combination with the DGK and 5-HT inhibitor, R-59-022. (B) Quantifications of GFP pixel intensity revealed that R-59-022, like Ritanserin, results in a synergistic reduction in tumour burden together with Trametinib. Data from 4 experiments, each with 16-24 replicate wells for each treatment. (C) Binarized image of a drug test plate where larvae were treated with Trametinib in combination with the DGK inhibitor, Amb639752. (D) Quantifications of GFP pixel intensity revealed that Amb639752, like Ritanserin, results in a synergistic reduction in tumour burden together with Trametinib. Data from 1 experiment with 24 replicate wells for each treatment. (E) Binarized image of a drug test plate where larvae were treated with Trametinib in combination with the DGK specific inhibitor, CU-3. (F) Quantifications of GFP pixel intensity revealed that CU-3, like Ritanserin, results in a synergistic reduction in tumour burden when combined with Trametinib. Data from 1 experiment with 24 replicate wells for each treatment. Statistical tests used were one-way ANOVAs with Tukey’s multiple comparisons. Error bars represent S.E.M. ** = p<0.01, *** = p<0.001, **** = p<0.0001.

**Supplementary Figure 5. Specific inhibition of serotonin receptors does not synergise with Trametinib to reduce *scrib^-^/Ras^V12^* tumour burden.**

(A, B) Binarized image of a drug test plate where larvae were treated with Trametinib in combination with the 5HT inhibitors, Paliperidone (A) or Volaninserin (B). (C, D) Quantifications of GFP pixel intensity revealed that Paliperidone or Volaninserin do not synergise with Trametinib to reduce tumour burden. Data from 1 experiment with 16 replicate wells for each treatment. Error bars represent S.E.M. A one-way ANOVA with a Tukey’s multiple comparison test was used to measure statistical significance. * p < 0.05, ** p < 0.01, *** p < 0.001, **** p < 0.0001.

**Supplementary Figure 6. RNAi lines utilised in this study are effective at reducing gene expression.**

Quantification of qRT-PCR data testing the efficacy of the various RNAi lines utilised in Figure 5 relative to the *luciferase-RNAi* non-targeting control. Gene expression measured in fold change normalised to the non-targeting RNAi control. All RNAi lines employed were found to be effective at reducing gene expression.

**Supplementary Figure 7 Trametinib synergises with Ritanserin in cultured MCF10A cells separately expressing either *shRNA-SCRIB* or human *H-RAS^G12V^*.**

(A) Western blotting confirmed SCRIB knockdown and H-RAS^G12V^ expression in MCF10A cells with the respective constructs. HSP70 was used as a loading control. (B,C) Dose response curves were generated for Trametinib (B) and Ritanserin (C) in MCF10A cells with knockdown of *SCRIB* or expression of *H-RAS^G12V^*. Live cell proportions were determined via CellTiter-Glo assay, and IC_50_ values were calculated from 3 independent experiments. (D,E) BLISS synergy scores for *H-RAS^G12V^* (D) and *shRNA-SCRIB* (E) MCF10A cells treated with Trametinib in combination with Ritanserin at the indicated doses. Doses approximate magnitudes of the IC_50_ for each drug. The proportion of live cells was determined by CellTiter-Glo assay, across 3 replicate experiments. A BLISS score > 0 indicates synergy.

## References

Abdel-Rahman, O., H. ElHalawani, and H. Ahmed. 2015. Risk of selected dermatological toxicities in cancer patients treated with MEK inhibitors: a comparative systematic review and meta-analysis. Future Oncol. 11:3307–3319.

Al-Jundi, M., S. Thakur, S. Gubbi, and J. Klubo-Gwiezdzinska. 2020. Novel Targeted Therapies for Metastatic Thyroid Cancer-A Comprehensive Review. Cancers (Basel). 12.

Andresen, B.T., M.A. Rizzo, K. Shome, and G. Romero. 2002. The role of phosphatidic acid in the regulation of the Ras/MEK/Erk signaling cascade. FEBS Letters. 531:65–68.

Anforth, R., M. Liu, B. Nguyen, P. Uribe, R. Kefford, A. Clements, G.V. Long, and P. Fernandez-Penas. 2014. Acneiform eruptions: a common cutaneous toxicity of the MEK inhibitor trametinib. Australas J Dermatol. 55:250–254.

Archibald, A., M. Al-Masri, A. Liew-Spilger, and L. McCaffrey. 2015. Atypical protein kinase C induces cell transformation by disrupting Hippo/Yap signaling. Mol Biol Cell. 26:3578–3595.

Arranz-Nicolas, J., and I. Merida. 2020. Biological regulation of diacylglycerol kinases in normal and neoplastic tissues: New opportunities for cancer immunotherapy. Adv Biol Regul. 75:100663.

Audia, A., and K.P. Bhat. 2018. Ritanserin, a novel agent targeting the mesenchymal subtype of glioblastomas. Neuro-Oncology. 20:151–152.

Bangi, E. 2019. A Drosophila Based Cancer Drug Discovery Framework. Adv Exp Med Biol. 1167:237–248.

Bangi, E., C. Ang, P. Smibert, A.V. Uzilov, A.G. Teague, Y. Antipin, R. Chen, C. Hecht, N. Gruszczynski, W.J. Yon, D. Malyshev, D. Laspina, I. Selkridge, H. Rainey, A.S. Moe, C.Y. Lau, P. Taik, E. Wilck, A. Bhardwaj, M. Sung, S. Kim, K. Yum, R. Sebra, M. Donovan, K. Misiukiewicz, E.E. Schadt, M.R. Posner, and R.L. Cagan. 2019. A personalized platform identifies trametinib plus zoledronate for a patient with KRAS-mutant metastatic colorectal cancer. Sci Adv. 5:eaav6528.

Bangi, E., D. Garza, and M. Hild. 2011. In vivo analysis of compound activity and mechanism of action using epistasis in Drosophila. Journal of Chemical Biology. 4:55–68.

Bangi, E., C. Murgia, A.G. Teague, O.J. Sansom, and R.L. Cagan. 2016. Functional exploration of colorectal cancer genomes using Drosophila. Nat Commun. 7:13615.

Bier, E. 2005. Drosophila, the golden bug, emerges as a tool for human genetics. Nature Reviews Genetics. 6:9.

Bilder, D. 2004. Epithelial polarity and proliferation control: links from the Drosophila neoplastic tumor suppressors. Genes & Development. 18:1909–1925.

Bliss, C.I. 1939. THE TOXICITY OF POISONS APPLIED JOINTLY. Annals of applied biology. 26:585–615.

Boroda, S., M. Niccum, V. Raje, B.W. Purow, and T.E. Harris. 2017. Dual activities of ritanserin and R59022 as DGKα inhibitors and serotonin receptor antagonists. Biochemical pharmacology. 123:29–39.

Bosco, E.E., J.C. Mulloy, and Y. Zheng. 2009. Rac1 GTPase: a “Rac” of all trades. Cell Mol Life Sci. 66:370–374.

Brumby, A.M., K.R. Goulding, T. Schlosser, S. Loi, R. Galea, P. Khoo, J.E. Bolden, T. Aigaki, P.O. Humbert, and H.E. Richardson. 2011. Identification of novel Ras-cooperating oncogenes in Drosophila melanogaster: a RhoGEF/Rho-family/JNK pathway is a central driver of tumorigenesis. Genetics. 188:105–125.

Brumby, A.M., and H.E. Richardson. 2003. scribble mutants cooperate with oncogenic Ras or Notch to cause neoplastic overgrowth in Drosophila. The EMBO Journal. 22:5769–5779.

Brumby, A.M., and H.E. Richardson. 2005. Using Drosophila melanogaster to map human cancer pathways. Nature Reviews Cancer. 5:626.

Carnero, A. 2010. The PKB/AKT pathway in cancer. Curr Pharm Des. 16:34–44.

Chen, J., W. Zhang, Y. Wang, D. Zhao, M. Wu, J. Fan, J. Li, Y. Gong, N. Dan, D. Yang, R. Liu, and Q. Zhan. 2019. The diacylglycerol kinase α (DGKα)/Akt/NF-κB feedforward loop promotes esophageal squamous cell carcinoma (ESCC) progression via FAK-dependent and FAK-independent manner. Oncogene. 38:2533-2550.

Chue, P., and J. Chue. 2012. A review of paliperidone palmitate. Expert Rev Neurother. 12:1383–1397.

Coleman, M.L., R.M. Densham, D.R. Croft, and M.F. Olson. 2006. Stability of p21Waf1/Cip1 CDK inhibitor protein is responsive to RhoA-mediated regulation of the actin cytoskeleton. Oncogene. 25:2708–2716.

Colon-Gonzalez, F., and M.G. Kazanietz. 2006. C1 domains exposed: from diacylglycerol binding to protein-protein interactions. Biochim Biophys Acta. 1761:827–837.

Cornish, J.W., I. Maany, P.J. Fudala, R.N. Ehrman, S.J. Robbins, and C.P. O’Brien. 2001. A randomized, double-blind, placebo-controlled study of ritanserin pharmacotherapy for cocaine dependence. Drug and Alcohol Dependence. 61:183–189.

Creed-Carson, M., A. Oraha, and J.N. Nobrega. 2011. Effects of 5-HT(2A) and 5-HT(2C) receptor antagonists on acute and chronic dyskinetic effects induced by haloperidol in rats. Behav Brain Res. 219:273–279.

Dar, A.C., T.K. Das, K.M. Shokat, and R.L. Cagan. 2012. Chemical genetic discovery of targets and anti-targets for cancer polypharmacology. Nature. 486:80.

De Leo, S., M. Trevisan, and L. Fugazzola. 2020. Recent advances in the management of anaplastic thyroid cancer. Thyroid Res. 13:17.

DeNicola, G.M., and D.A. Tuveson. 2009. RAS in cellular transformation and senescence. Eur J Cancer. 45 Suppl 1:211–216.

Dimauro, T., and G. David. 2010. Ras-induced senescence and its physiological relevance in cancer. Curr Cancer Drug Targets. 10:869–876.

Doggett, K., F.A. Grusche, H.E. Richardson, and A.M. Brumby. 2011. Loss of the Drosophila cell polarity regulator Scribbled promotes epithelial tissue overgrowth and cooperation with oncogenic Ras-Raf through impaired Hippo pathway signaling. BMC Developmental Biology. 11:57.

Dominguez, C.L., D.H. Floyd, A. Xiao, G.R. Mullins, B.A. Kefas, W. Xin, M.N. Yacur, R. Abounader, J.K. Lee, G.M. Wilson, T.E. Harris, and B.W. Purow. 2013. Diacylglycerol kinase alpha is a critical signaling node and novel therapeutic target in glioblastoma and other cancers. Cancer Discov. 3:782–797.

Dow, L.E., I.A. Elsum, C.L. King, K.M. Kinross, H.E. Richardson, and P.O. Humbert. 2008. Loss of human Scribble cooperates with H-Ras to promote cell invasion through deregulation of MAPK signalling. Oncogene. 27:5988.

Ebdrup, B.H., H. Rasmussen, J. Arnt, and B. Glenthoj. 2011. Serotonin 2A receptor antagonists for treatment of schizophrenia. Expert Opin Investig Drugs. 20:1211–1223.

Edwards, A., M. Gladstone, P. Yoon, D. Raben, B. Frederick, and T.T. Su. 2011. Combinatorial effect of maytansinol and radiation in *Drosophila* and human cancer cells. Disease Models & Mechanisms. 4:496–503.

Elsum, I., L. Yates, P.O. Humbert, and H.E. Richardson. 2012. The Scribble–Dlg–Lgl polarity module in development and cancer: from flies to man. Essays In Biochemistry. 53:141–168.

Fang, Y., M. Vilella-Bach, R. Bachmann, A. Flanigan, and J. Chen. 2001. Phosphatidic acid-mediated mitogenic activation of mTOR signaling. Science. 294:1942–1945.

Fazio, A., E. Owusu Obeng, I. Rusciano, M.V. Marvi, M. Zoli, S. Mongiorgi, G. Ramazzotti, M.Y. Follo, J.A. McCubrey, L. Cocco, L. Manzoli, and S. Ratti. 2020. Subcellular Localization Relevance and Cancer-Associated Mechanisms of Diacylglycerol Kinases. Int J Mol Sci. 21.

Feigin, M.E., S.D. Akshinthala, K. Araki, A.Z. Rosenberg, L.B. Muthuswamy, B. Martin, B.D. Lehmann, H.K. Berman, J.A. Pietenpol, R.D. Cardiff, and S.K. Muthuswamy. 2014. Mislocalization of the Cell Polarity Protein Scribble Promotes Mammary Tumorigenesis and Is Associated with Basal Breast Cancer. Cancer Research. 74:3180.

Ferlay, J., I. Soerjomataram, R. Dikshit, S. Eser, C. Mathers, M. Rebelo, D.M. Parkin, D. Forman, and F. Bray. 2015. Cancer incidence and mortality worldwide: Sources, methods and major patterns in GLOBOCAN 2012. International Journal of Cancer. 136:E359–E386.

Ferrari, S.M., G. Elia, F. Ragusa, I. Ruffilli, C. La Motta, S.R. Paparo, A. Patrizio, R. Vita, S. Benvenga, G. Materazzi, P. Fallahi, and A. Antonelli. 2020. Novel treatments for anaplastic thyroid carcinoma. Gland Surg. 9:S28–S42.

Foster, D.A., D. Salloum, D. Menon, and M.A. Frias. 2014. Phospholipase D and the maintenance of phosphatidic acid levels for regulation of mammalian target of rapamycin (mTOR). J Biol Chem. 289:22583–22588.

Franks, C.E., S.T. Campbell, B.W. Purow, T.E. Harris, and K.-L. Hsu. 2017. The Ligand Binding Landscape of Diacylglycerol Kinases. Cell Chemical Biology. 24:870–880.e875.

Froldi, F., M. Ziosi, G. Tomba, F. Parisi, F. Garoia, A. Pession, and D. Grifoni. 2008. Drosophila Lethal Giant Larvae Neoplastic Mutant as a Genetic Tool for Cancer Modeling. Current Genomics. 9:147–154.

Gjertsen, B.T., and P. Schöffski. 2014. Discovery and development of the Polo-like kinase inhibitor volasertib in cancer therapy. Leukemia. 29:11.

Gladstone, M., B. Frederick, D. Zheng, A. Edwards, P. Yoon, S. Stickel, T. DeLaney, D.C. Chan, D. Raben, and T.T. Su. 2012. A translation inhibitor identified in a *Drosophila* screen enhances the effect of ionizing radiation and taxol in mammalian models of cancer. Disease Models &; Mechanisms. 5:342–350.

Gladstone, M., and T.T. Su. 2011. Chemical genetics and drug screening in Drosophila cancer models. Journal of Genetics and Genomics. 38:497–504.

Godde, N.J., H.B. Pearson, L.K. Smith, and P.O. Humbert. 2014. Dissecting the role of polarity regulators in cancer through the use of mouse models. Exp Cell Res. 328:249–257.

Goldoni, M., and S. Tagliaferri. 2011. Dose-response or dose-effect curves in in vitro experiments and their use to study combined effects of neurotoxicants. Methods Mol Biol. 758:415–434.

Gonzalez, C. 2013. Drosophila melanogaster: a model and a tool to investigate malignancy and identify new therapeutics. Nature Reviews Cancer. 13:172.

Griner, E.M., and M.G. Kazanietz. 2007. Protein kinase C and other diacylglycerol effectors in cancer. Nat Rev Cancer. 7:281–294.

Grzeschik, N.A., L.M. Parsons, M.L. Allott, K.F. Harvey, and H.E. Richardson. 2010. Lgl, aPKC, and Crumbs regulate the Salvador/Warts/Hippo pathway through two distinct mechanisms. Curr Biol. 20:573–581.

Hanahan, D., and R.A. Weinberg. 2011. Hallmarks of Cancer: The Next Generation. Cell. 144:646–674.

Harrington, L.S., G.M. Findlay, and R.F. Lamb. 2005. Restraining PI3K: mTOR signalling goes back to the membrane. Trends Biochem Sci. 30:35–42.

Hay, M., D.W. Thomas, J.L. Craighead, C. Economides, and J. Rosenthal. 2014. Clinical development success rates for investigational drugs. Nature Biotechnology. 32:40.

Hedner, T., and B. Persson. 1988. Effects of a new serotonin antagonist, ketanserin, in experimental and clinical hypertension. Am J Hypertens. 1:317S-323S.

Hoffner, B., and K. Benchich. 2018. Trametinib: A Targeted Therapy in Metastatic Melanoma. J Adv Pract Oncol. 9:741–745.

Humbert, P.O., N.A. Grzeschik, A.M. Brumby, R. Galea, I. Elsum, and H.E. Richardson. 2008. Control of tumourigenesis by the Scribble/Dlg/Lgl polarity module. Oncogene. 27:6888.

Huser, A., M. Eschment, N. Güllü, K.A.N. Collins, K. Böpple, L. Pankevych, E. Rolsing, and A.S. Thum. 2017. Anatomy and behavioral function of serotonin receptors in Drosophila melanogaster larvae. PLOS ONE. 12:e0181865.

Igaki, T., R.A. Pagliarini, and T. Xu. 2006. Loss of Cell Polarity Drives Tumor Growth and Invasion through JNK Activation in *Drosophila*. Current Biology. 16:1139–1146.

Ito, T., and T. Igaki. 2021. Yorkie drives Ras-induced tumor progression by microRNA-mediated inhibition of cellular senescence. Sci Signal. 14.

Jaklevic, B., L. Uyetake, W. Lemstra, J. Chang, W. Leary, A. Edwards, S. Vidwans, O. Sibon, and T. Tin Su. 2006. Contribution of Growth and Cell Cycle Checkpoints to Radiation Survival in Drosophila. Genetics. 174:1963–1972.

Johnson, B.A., D.R. Jasinski, G.P. Galloway, H. Kranzler, R. Weinreib, R.F. Anton, B.J. Mason, M.J. Bohn, H.M. Pettinati, R. Rawson, C. Clyde, and G. Ritanserin Study. 1996. Ritanserin in the treatment of alcohol dependence – a multi-center clinical trial. Psychopharmacology. 128:206–215.

Jones, M.T., M.T. Strassnig, and P.D. Harvey. 2020. Emerging 5-HT receptor antagonists for the treatment of Schizophrenia. Expert Opin Emerg Drugs. 25:189–200.

Kai, M., S. Yasuda, S. Imai, M. Toyota, H. Kanoh, and F. Sakane. 2009. Diacylglycerol kinase alpha enhances protein kinase Czeta-dependent phosphorylation at Ser311 of p65/RelA subunit of nuclear factor-kappaB. FEBS Lett. 583:3265–3268.

Karim, F.D., and G.M. Rubin. 1998. Ectopic expression of activated Ras1 induces hyperplastic growth and increased cell death in Drosophila imaginal tissues. Development. 125:1–9.

La Marca, J.E., S.T. Diepstraten, A.L. Hodge, H. Wang, A.H. Hart, H.E. Richardson, and W.G. Somers. 2019. Strip and Cka negatively regulate JNK signalling during Drosophila spermatogenesis. Development. 146.

Lee, M., and V. Vasioukhin. 2008. Cell polarity and cancer--cell and tissue polarity as a non-canonical tumor suppressor. J Cell Sci. 121:1141–1150.

Leong, G.R., K.R. Goulding, N. Amin, H.E. Richardson, and A.M. Brumby. 2009. *scribble* mutants promote aPKC and JNK-dependent epithelial neoplasia independently of Crumbs. BMC Biology. 7:62.

Levine, B.D., and R.L. Cagan. 2016. Drosophila Lung Cancer Models Identify Trametinib plus Statin as Candidate Therapeutic. Cell Rep. 14:1477–1487.

Leysen, J.E., W. Gommeren, P. Van Gompel, J. Wynants, P.F. Janssen, and P.M. Laduron. 1985. Receptor-binding properties in vitro and in vivo of ritanserin: A very potent and long acting serotonin-S2 antagonist. Mol Pharmacol. 27:600–611.

Li, X., H. Yang, J. Liu, M.D. Schmidt, and T. Gao. 2011. Scribble-mediated membrane targeting of PHLPP1 is required for its negative regulation of Akt. EMBO Rep. 12:818–824.

Liu, K., N. Kunii, M. Sakuma, A. Yamaki, S. Mizuno, M. Sato, H. Sakai, S. Kado, K. Kumagai, H. Kojima, T. Okabe, T. Nagano, Y. Shirai, and F. Sakane. 2016. A novel diacylglycerol kinase α-selective inhibitor, CU-3, induces cancer cell apoptosis and enhances immune response. Journal of Lipid Research. 57:368-379.

Ma, X., W. Xu, D. Zhang, Y. Yang, W. Li, and L. Xue. 2015. Wallenda regulates JNK-mediated cell death in Drosophila. Cell Death & Disease. 6:e1737.

Malumbres, M., and M. Barbacid. 2003. RAS oncogenes: the first 30 years. Nat Rev Cancer. 3:459–465.

Manning, B.D., M.N. Logsdon, A.I. Lipovsky, D. Abbott, D.J. Kwiatkowski, and L.C. Cantley. 2005. Feedback inhibition of Akt signaling limits the growth of tumors lacking Tsc2. Genes Dev. 19:1773–1778.

Manoharan, N., J. Choi, C. Chordas, M.A. Zimmerman, J. Scully, J. Clymer, M. Filbin, N.J. Ullrich, P. Bandopadhayay, S.N. Chi, and K.K. Yeo. 2020. Trametinib for the treatment of recurrent/progressive pediatric low-grade glioma. J Neurooncol. 149:253–262.

Markstein, M., S. Dettorre, J. Cho, R.A. Neumüller, S. Craig-Müller, and N. Perrimon. 2014. Systematic screen of chemotherapeutics in *Drosophila* stem cell tumors. Proceedings of the National Academy of Sciences. 111:4530–4535.

Masahiro, S., P.S. Alex, M.U.U. Peter, A.M. Matthew, S. Lisa, Y.M. Andres, R. Alexander, S. Avner, L.C. Ross, and C.D. Arvin. 2018. A whole-animal platform to advance a clinical kinase inhibitor into new disease space. Nature Chemical Biology. 14.

Memmott, R.M., and P.A. Dennis. 2009. Akt-dependent and -independent mechanisms of mTOR regulation in cancer. Cell Signal. 21:656–664.

Merida, I., A. Avila-Flores, and E. Merino. 2008. Diacylglycerol kinases: at the hub of cell signalling. Biochem J. 409:1–18.

Mérida, I., A. Ávila-Flores, and E. Merino. 2008. Diacylglycerol kinases: at the hub of cell signalling. Biochemical Journal. 409:1–18.

Merida, I., P. Torres-Ayuso, A. Avila-Flores, J. Arranz-Nicolas, E. Andrada, M. Tello-Lafoz, R. Liebana, and R. Arcos. 2017. Diacylglycerol kinases in cancer. Adv Biol Regul. 63:22–31.

Mizutani, K., S. Sonoda, and H. Wakita. 2018. Ritanserin, a serotonin-2 receptor antagonist, inhibits functional recovery after cerebral infarction. NeuroReport. 29:54–58.

Muthuswamy, S.K., and B. Xue. 2012. Cell polarity as a regulator of cancer cell behavior plasticity. Annu Rev Cell Dev Biol. 28:599–625.

Mutterer, J., and W. Rasband. 2012. ImageJ Macro Language -Programmer’s Reference Guide. Vol. v1.46d. p1-45. https://imagej.nih.gov/ij/docs/macro_reference_guide.pdf.

Nagasaka, K., D. Pim, P. Massimi, M. Thomas, V. Tomaic, V.K. Subbaiah, C. Kranjec, S. Nakagawa, T. Yano, Y. Taketani, M. Myers, and L. Banks. 2010. The cell polarity regulator hScrib controls ERK activation through a KIM site-dependent interaction. Oncogene. 29:5311–5321.

Nagasaka, K., T. Seiki, A. Yamashita, P. Massimi, V.K. Subbaiah, M. Thomas, C. Kranjec, K. Kawana, S. Nakagawa, T. Yano, Y. Taketani, T. Fujii, S. Kozuma, and L. Banks. 2013. A novel interaction between hScrib and PP1gamma downregulates ERK signaling and suppresses oncogene-induced cell transformation. PLoS One. 8:e53752.

Nakamura, M., and T. Igaki. 2017. Induction and Detection of Oncogene-Induced Cellular Senescence in Drosophila. Methods Mol Biol. 1534:211–218.

Nappi, G., G. Sandrini, F. Granella, L. Ruiz, G. Cerutti, F. Facchinetti, F. Blandini, and G.C. Manzoni. 1990. A new 5-HT2 antagonist (ritanserin) in the treatment of chronic headache with depression. A double-blind study vs amitriptyline. Headache. 30:439–444.

Norman, M., K.A. Wisniewska, K. Lawrenson, P. Garcia-Miranda, M. Tada, M. Kajita, H. Mano, S. Ishikawa, M. Ikegawa, T. Shimada, and Y. Fujita. 2012. Loss of Scribble causes cell competition in mammalian cells. J Cell Sci. 125:59–66.

Oliva, J.L., E.M. Griner, and M.G. Kazanietz. 2005. PKC isozymes and diacylglycerol-regulated proteins as effectors of growth factor receptors. Growth Factors. 23:245–252.

Olmez, I., S. Love, A. Xiao, L. Manigat, P. Randolph, B.D. McKenna, B.P. Neal, S. Boroda, M. Li, B. Brenneman, R. Abounader, D. Floyd, J. Lee, I. Nakano, J. Godlewski, A. Bronisz, E.P. Sulman, M. Mayo, D. Gioeli, M. Weber, T.E. Harris, and B. Purow. 2018. Targeting the mesenchymal subtype in glioblastoma and other cancers via inhibition of diacylglycerol kinase alpha. Neuro-Oncology. 20:192–202.

Olson, M.F., H.F. Paterson, and C.J. Marshall. 1998. Signals from Ras and Rho GTPases interact to regulate expression of p21Waf1/Cip1. Nature. 394:295–299.

Pagliarini, R.A., and T. Xu. 2003. A Genetic Screen in *Drosophila* for Metastatic Behavior. Science. 302:1227–1231.

Pearson, H.B., P.A. Perez-Mancera, L.E. Dow, A. Ryan, P. Tennstedt, D. Bogani, I. Elsum, A. Greenfield, D.A. Tuveson, R. Simon, and P.O. Humbert. 2011. SCRIB expression is deregulated in human prostate cancer, and its deficiency in mice promotes prostate neoplasia. J Clin Invest. 121:4257–4267.

Posch, C., B.D. Cholewa, I. Vujic, M. Sanlorenzo, J. Ma, S.T. Kim, S. Kleffel, T. Schatton, K. Rappersberger, R. Gutteridge, N. Ahmad, and S. Ortiz/Urda. 2015. Combined inhibition of MEK and Plk1 has synergistic anti-tumor activity in NRAS mutant melanoma. The Journal of investigative dermatology. 135:2475–2483.

Pylayeva-Gupta, Y., E. Grabocka, and D. Bar-Sagi. 2011. RAS oncogenes: weaving a tumorigenic web. Nat Rev Cancer. 11:761–774.

Ratheesh, A., R. Priya, and A.S. Yap. 2013. Coordinating Rho and Rac: the regulation of Rho GTPase signaling and cadherin junctions. Prog Mol Biol Transl Sci. 116:49–68.

Richardson, H.E., L. Willoughby, and P.O. Humbert. 2015. Screening for Anti-cancer Drugs in Drosophila. *In* eLS. John Wiley & Sons, Ltd.

Roskoski, R., Jr. 2017. Allosteric MEK1/2 inhibitors including cobimetanib and trametinib in the treatment of cutaneous melanomas. Pharmacol Res. 117:20–31.

Royer, C., and X. Lu. 2011. Epithelial cell polarity: a major gatekeeper against cancer? Cell Death Differ. 18:1470–1477.

Rudrapatna, V.A., R.L. Cagan, and T.K. Das. 2012. Drosophila cancer models. Dev Dyn. 241:107–118.

Sahai, E., M.F. Olson, and C.J. Marshall. 2001. Cross-talk between Ras and Rho signalling pathways in transformation favours proliferation and increased motility. EMBO J. 20:755–766.

Sakane, F., F. Hoshino, M. Ebina, H. Sakai, and D. Takahashi. 2021. The Roles of Diacylglycerol Kinase alpha in Cancer Cell Proliferation and Apoptosis. Cancers (Basel). 13.

Sakane, F., S. Imai, M. Kai, S. Yasuda, and H. Kanoh. 2008. Diacylglycerol kinases as emerging potential drug targets for a variety of diseases. Curr Drug Targets. 9:626–640.

Sakane, F., S. Mizuno, and S. Komenoi. 2016. Diacylglycerol Kinases as Emerging Potential Drug Targets for a Variety of Diseases: An Update. Front Cell Dev Biol. 4:82.

Sato, M., K. Liu, S. Sasaki, N. Kunii, H. Sakai, H. Mizuno, H. Saga, and F. Sakane. 2013. Evaluations of the Selectivities of the Diacylglycerol Kinase Inhibitors R59022 and R59949 Among Diacylglycerol Kinase Isozymes Using a New Non-Radioactive Assay Method. Pharmacology. 92:99–107.

Saxton, R.A., and D.M. Sabatini. 2017. mTOR Signaling in Growth, Metabolism, and Disease. Cell. 168:960–976.

Schindelin, J., I. Arganda-Carreras, E. Frise, V. Kaynig, M. Longair, T. Pietzsch, S. Preibisch, C. Rueden, S. Saalfeld, and B. Schmid. 2012. Fiji: an open-source platform for biological-image analysis. Nature methods. 9:676.

Schoffski, P. 2009. Polo-like kinase (PLK) inhibitors in preclinical and early clinical development in oncology. Oncologist. 14:559–570.

Shakeel, I., N. Basheer, G.M. Hasan, M. Afzal, and M.I. Hassan. 2021. Polo-like Kinase 1 as an emerging drug target: structure, function and therapeutic implications. J Drug Target. 29:168–184.

Siegel, R.L., K.D. Miller, and A. Jemal. 2016. Cancer statistics, 2016. CA: A Cancer Journal for Clinicians. 66:7-30.

Sonoshita, M., and R.L. Cagan. 2017. Chapter Nine - Modeling Human Cancers in Drosophila. *In* Current Topics in Developmental Biology. Vol. 121. L. Pick, editor. Academic Press. 287–309.

Stark, M.B. 1918. An Hereditary Tumor in the Fruit Fly, Drosophila. The Journal of Cancer Research. 3:279–301.

Stephens, R., K. Lim, M. Portela, M. Kvansakul, P.O. Humbert, and H.E. Richardson. 2018. The Scribble Cell Polarity Module in the Regulation of Cell Signaling in Tissue Development and Tumorigenesis. J Mol Biol. 430:3585–3612.

Stickel, S., N. Gomes, B. Frederick, D. Raben, and T. Su. 2015. Bouvardin is a Radiation Modulator with a Novel Mechanism of Action. Radiation Research. 184:392–403.

Tabbo, F., C. Pisano, J. Mazieres, L. Mezquita, E. Nadal, D. Planchard, A. Pradines, D. Santamaria, A. Swalduz, C. Ambrogio, S. Novello, S. Ortiz-Cuaran, and B. Consortium. 2022. How far we have come targeting BRAF-mutant non-small cell lung cancer (NSCLC). Cancer Treat Rev. 103:102335.

Tepass, U. 2012. The apical polarity protein network in Drosophila epithelial cells: regulation of polarity, junctions, morphogenesis, cell growth, and survival. Annu Rev Cell Dev Biol. 28:655–685.

Thomas, M., N. Narayan, D. Pim, V. Tomaić, P. Massimi, K. Nagasaka, C. Kranjec, N. Gammoh, and L. Banks. 2008. Human papillomaviruses, cervical cancer and cell polarity. Oncogene. 27:7018.

Toschi, A., E. Lee, L. Xu, A. Garcia, N. Gadir, and D.A. Foster. 2009. Regulation of mTORC1 and mTORC2 complex assembly by phosphatidic acid: competition with rapamycin. Mol Cell Biol. 29:1411–1420.

Uhlirova, M., and D. Bohmann. 2006. JNK- and Fos-regulated Mmp1 expression cooperates with Ras to induce invasive tumors in *Drosophila*. The EMBO Journal. 25:5294.

Velnati, S., S. Centonze, F. Girivetto, and G. Baldanzi. 2021a. Diacylglycerol Kinase alpha in X Linked Lymphoproliferative Disease Type 1. Int J Mol Sci. 22.

Velnati, S., S. Centonze, F. Girivetto, D. Capello, R.M. Biondi, A. Bertoni, R. Cantello, B. Ragnoli, M. Malerba, A. Graziani, and G. Baldanzi. 2021b. Identification of Key Phospholipids That Bind and Activate Atypical PKCs. Biomedicines. 9.

Velnati, S., A. Massarotti, A. Antona, M. Talmon, L.G. Fresu, A.S. Galetto, D. Capello, A. Bertoni, V. Mercalli, A. Graziani, G.C. Tron, and G. Baldanzi. 2020. Structure activity relationship studies on Amb639752: toward the identification of a common pharmacophoric structure for DGKalpha inhibitors. J Enzyme Inhib Med Chem. 35:96–108.

Velnati, S., E. Ruffo, A. Massarotti, M. Talmon, K.S.S. Varma, A. Gesu, L.G. Fresu, A.L. Snow, A. Bertoni, D. Capello, G.C. Tron, A. Graziani, and G. Baldanzi. 2019. Identification of a novel DGKα inhibitor for XLP-1 therapy by virtual screening. European Journal of Medicinal Chemistry. 164:378–390.

Vidal, M., S. Wells, A. Ryan, and R. Cagan. 2005. ZD6474 Suppresses Oncogenic RET Isoforms in a *Drosophila* Model for Type 2 Multiple Endocrine Neoplasia Syndromes and Papillary Thyroid Carcinoma. Cancer Research. 65:3538–3541.

Vorhagen, S., and C.M. Niessen. 2014. Mammalian aPKC/Par polarity complex mediated regulation of epithelial division orientation and cell fate. Exp Cell Res. 328:296–302.

Wang, H., and M.G. Kazanietz. 2006. The lipid second messenger diacylglycerol as a negative regulator of Rac signalling. Biochem Soc Trans. 34:855–857.

Wangler, M.F., S. Yamamoto, and H.J. Bellen. 2015. Fruit Flies in Biomedical Research. Genetics. 199:639–653.

Wangler, M.F., S. Yamamoto, H.-T. Chao, J.E. Posey, M. Westerfield, J. Postlethwait, P. Hieter, K.M. Boycott, P.M. Campeau, and H.J. Bellen. 2017. Model Organisms Facilitate Rare Disease Diagnosis and Therapeutic Research. Genetics. 207:9–27.

Wiesel, F.-A., A.L. Nordström, L. Farde, and B. Eriksson. 1994. An open clinical and biochemical study of ritanserin in acute patients with schizophrenia. Psychopharmacology. 114:31–38.

Willecke, M., J. Toggweiler, and K. Basler. 2011. Loss of PI3K blocks cell-cycle progression in a Drosophila tumor model. Oncogene. 30:4067–4074.

Willoughby, L.F., T. Schlosser, S.A. Manning, J.P. Parisot, I.P. Street, H.E. Richardson, P.O. Humbert, and A.M. Brumby. 2013. An *in vivo* large-scale chemical screening platform using *Drosophila* for anti-cancer drug discovery. Disease Models & Mechanisms. 6:521–529.

Wright, C.J., and P.L. McCormack. 2013. Trametinib: first global approval. Drugs. 73:1245–1254.

Yadav, A.K., S. Srikrishna, and S.C. Gupta. 2016. Cancer Drug Development Using Drosophila as an in vivo Tool: From Bedside to Bench and Back. Trends in Pharmacological Sciences. 37:789–806.

Yamaki, A., R. Akiyama, C. Murakami, S. Takao, Y. Murakami, S. Mizuno, D. Takahashi, S. Kado, A. Taketomi, Y. Shirai, K. Goto, and F. Sakane. 2019. Diacylglycerol kinase α-selective inhibitors induce apoptosis and reduce viability of melanoma and several other cancer cell lines. Journal of Cellular Biochemistry. 120:10043–10056.

Zeiser, R. 2014. Trametinib. Recent Results Cancer Res. 201:241–248.

Zhan, L., A. Rosenberg, K.C. Bergami, M. Yu, Z. Xuan, A.B. Jaffe, C. Allred, and S.K. Muthuswamy. 2008. Deregulation of Scribble Promotes Mammary Tumorigenesis and Reveals a Role for Cell Polarity in Carcinoma. Cell. 135:865–878.

Zhang, X., C. Wei, H. Liang, and L. Han. 2021. Polo-Like Kinase 4’s Critical Role in Cancer Development and Strategies for Plk4-Targeted Therapy. Front Oncol. 11:587554.

Zhang, Y., and G. Du. 2009. Phosphatidic acid signaling regulation of Ras superfamily of small guanosine triphosphatases. Biochim Biophys Acta. 1791:850–855.

