## Supplementary material for "A *Drosophila in vivo* chemical screen reveals that combination drug treatment targeting MEK and DGKα mitigates Ras-driven polarity-impaired tumourigenesis": Supp File 2

**Supplementary File** – Hits from the primary drug screen that did not significantly reduce *scrib⁻ Ras$^{V12}$* tumour burden upon retesting.

---

**Compound:** AC-93253 iodide
**Reported Function:** Cell permeable, selective RAR agonist, Src inhibitor

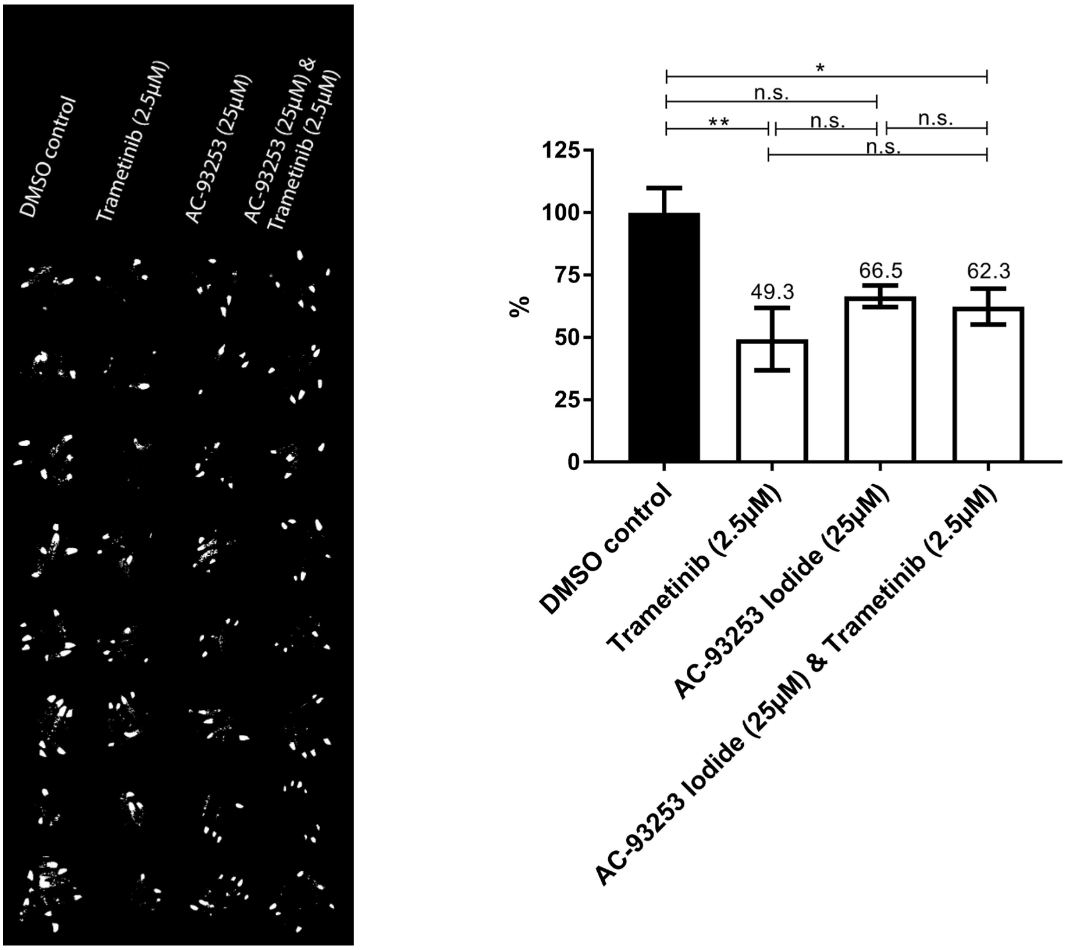


**Left** – Partial binarized image of screening plate showing various AC-932532 Iodide treatments. **Right** – Mean GFP pixel area, represented as a percentage of mean DMSO pixel area, for various AC-93253 Iodide treatments. N = 8 replicate wells per treatment. Error bars represent S.E.M. A one-way ANOVA, with a Tukey's multiple comparison test, was used to measure statistical significance. * $p < 0.05$, ** $p < 0.01$.

**Compound:** Carbinoxamine Maleate Salt
**Reported Function:** Histamine H1 receptor antagonist

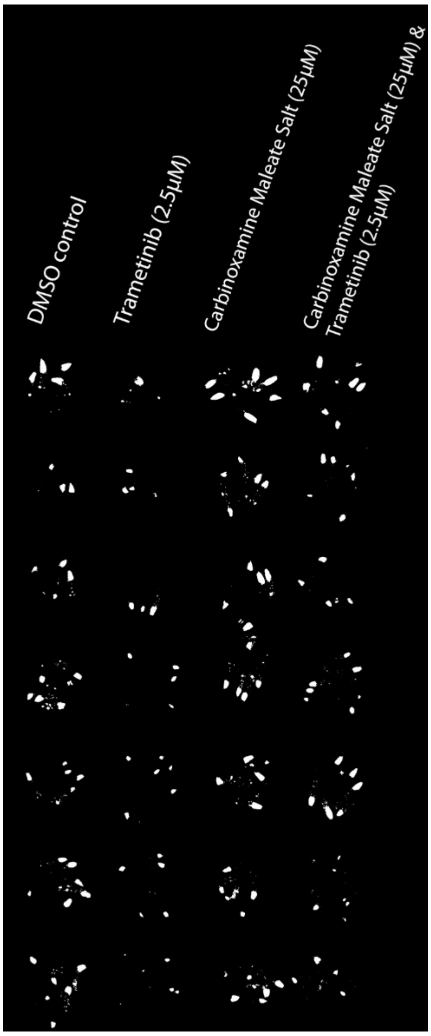


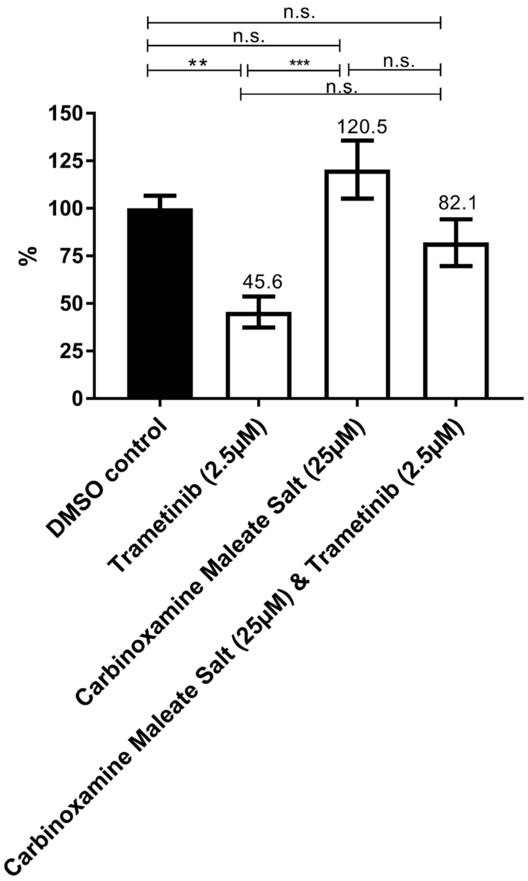


**Left** – Partial binarized image of screening plate showing various Carbinoxamine Maleate Salt treatments. **Right** – Mean GFP pixel area, represented as a percentage of mean DMSO pixel area, for various Carbinoxamine Maleate Salt treatments. N = 8 replicate wells per treatment. Error bars represent S.E.M. A one-way ANOVA, with a Tukey's multiple comparison test, was used to measure statistical significance. ** $p < 0.01$, *** $p < 0.001$.

**Compound:** CGP 57380
**Reported Function:** selective inhibitor of MNK1

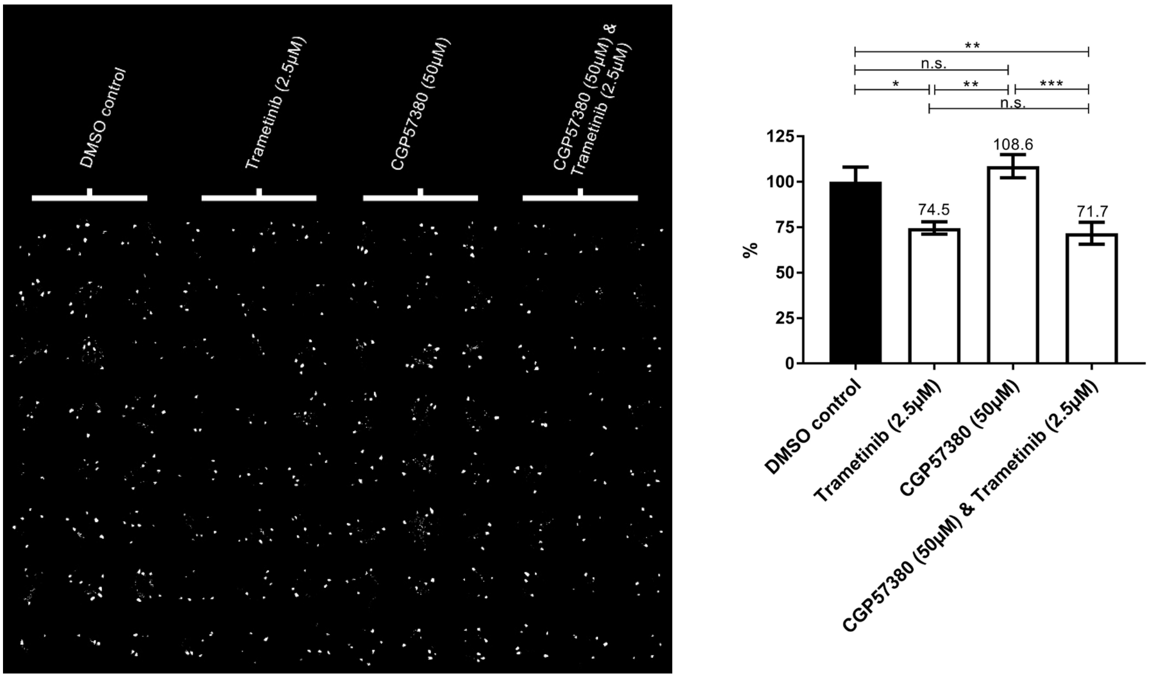


**Left** – Binarized image of screening plate showing various CGP57380 treatments. **Right** – Mean GFP pixel area, represented as a percentage of mean DMSO pixel area, for various CGP57380 treatments. N = 24 replicate wells per treatment. Error bars represent S.E.M. A one-way ANOVA, with a Tukey's multiple comparison test, was used to measure statistical significance. * $p < 0.05$, ** $p < 0.01$, *** $p < 0.001$.

**Compound:** Guvacine (hydrochloride)
**Reported Function:** Amino acid found in *Areca catechu*. Inhibits GABA uptake

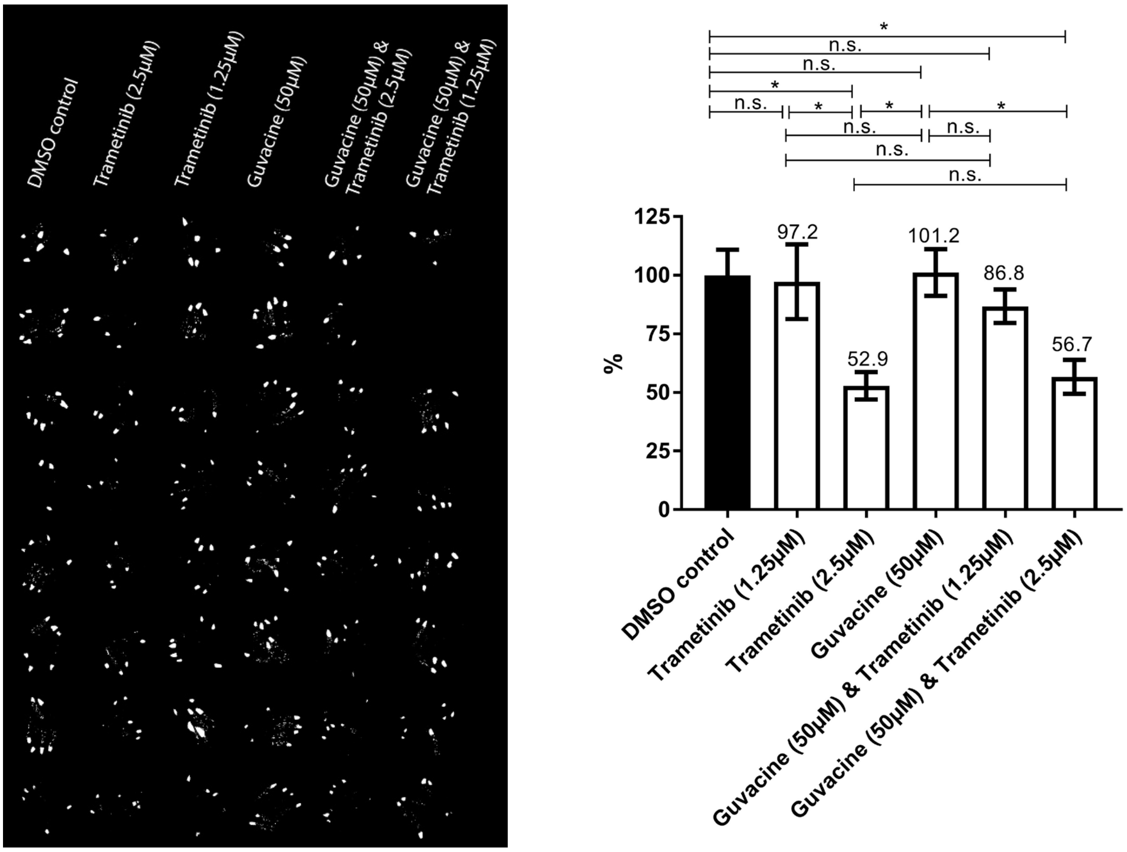


**Left** – Partial binarized image of screening plate showing various Guvacine treatments. **Right** – Mean GFP pixel area, represented as a percentage of mean DMSO pixel area, for various Guvacine treatments. N = 8 replicate wells per treatment (*N=7 replicate wells for Guvacine 50μM & Trametinib 2.5μM treatment*). Error bars represent S.E.M. A one-way ANOVA, with a Tukey's multiple comparison test, was used to measure statistical significance. * $p < 0.05$.

**Compound:** Hesperidin
**Reported Function:** Bioflavinoid

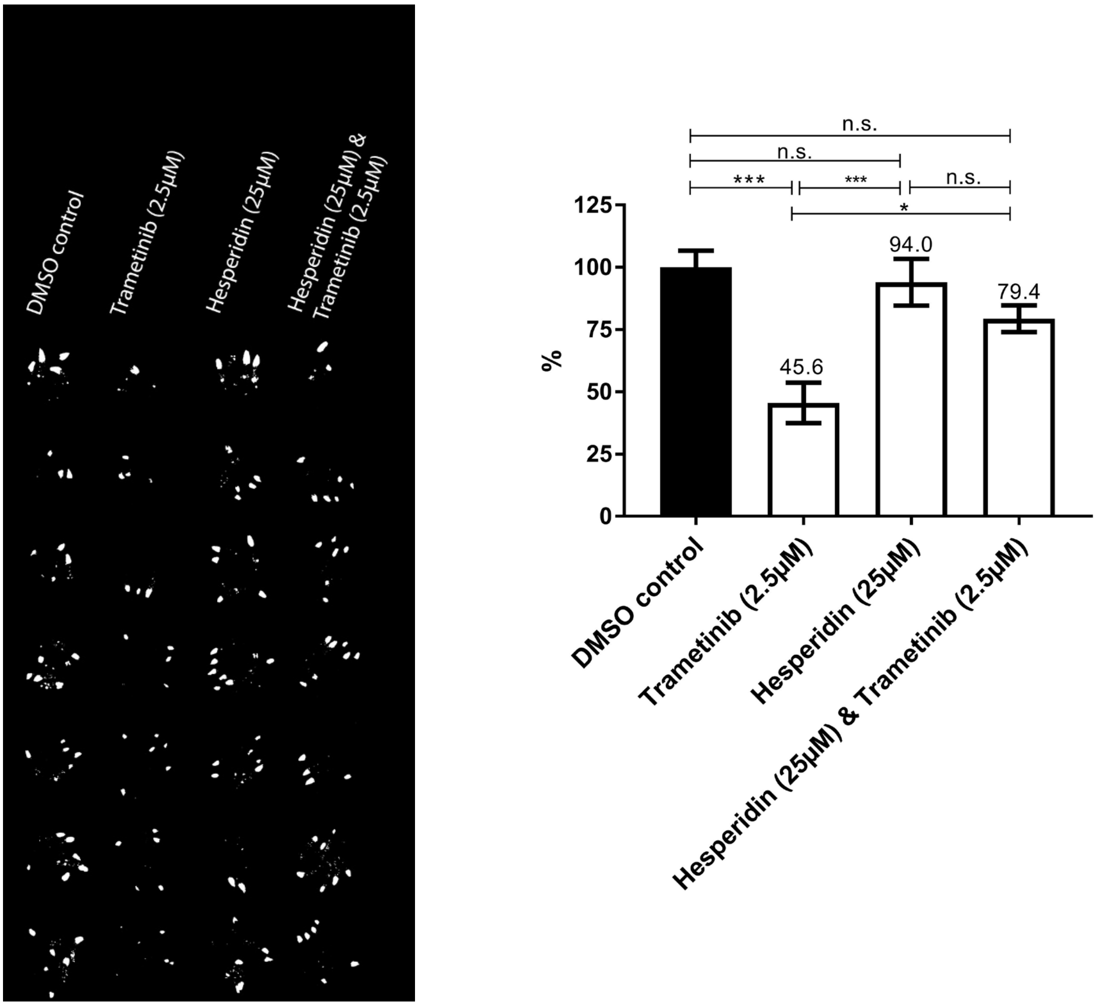


**Left** – Partial binarized image of screening plate showing various Hesperidin treatments. **Right** – Mean GFP pixel area, represented as a percentage of mean DMSO pixel area, for various Hesperidin treatments. N = 8 replicate wells per treatment. Error bars represent S.E.M. A one-way ANOVA, with a Tukey's multiple comparison test, was used to measure statistical significance. * $p < 0.05$, *** $p < 0.001$.

**Compound:** JS-K
**Reported Function:** Nitric oxide donor.

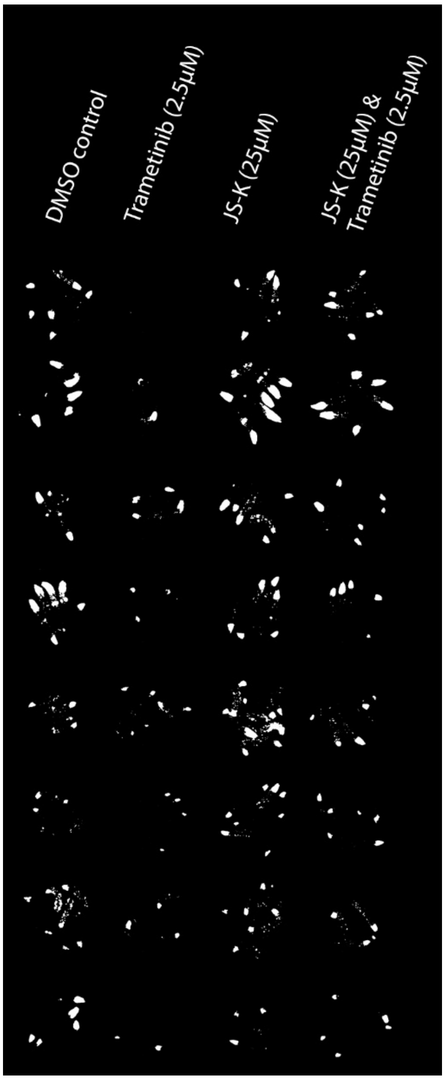


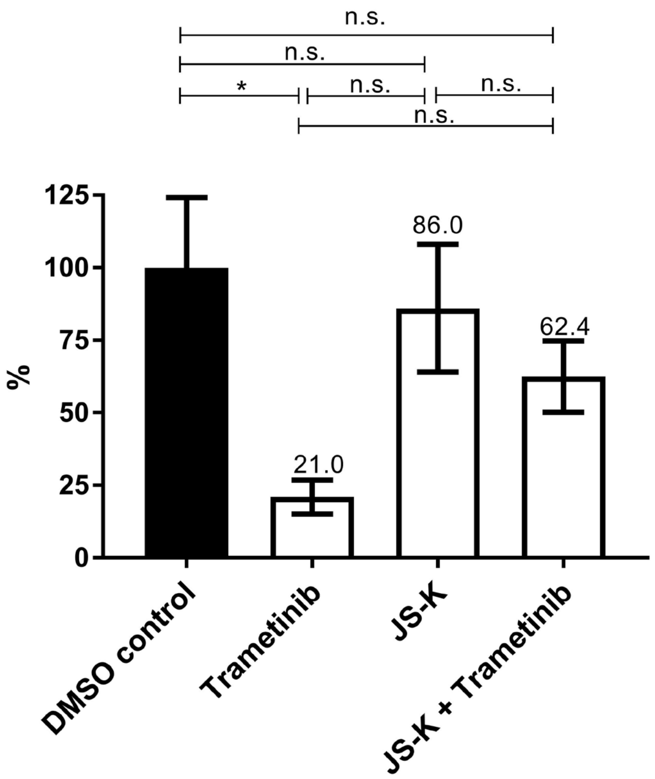


**Left** – Partial binarized image of screening plate showing various JS-K treatments. **Right** – Mean GFP pixel area, represented as a percentage of mean DMSO pixel area, for various JS-K treatments. N = 8 replicate wells per treatment. Error bars represent S.E.M. A one-way ANOVA, with a Tukey's multiple comparison test, was used to measure statistical significance. * $p < 0.05$.

**Compound:** LY294002
**Reported Function:** Broad spectrum PI3K inhibitor

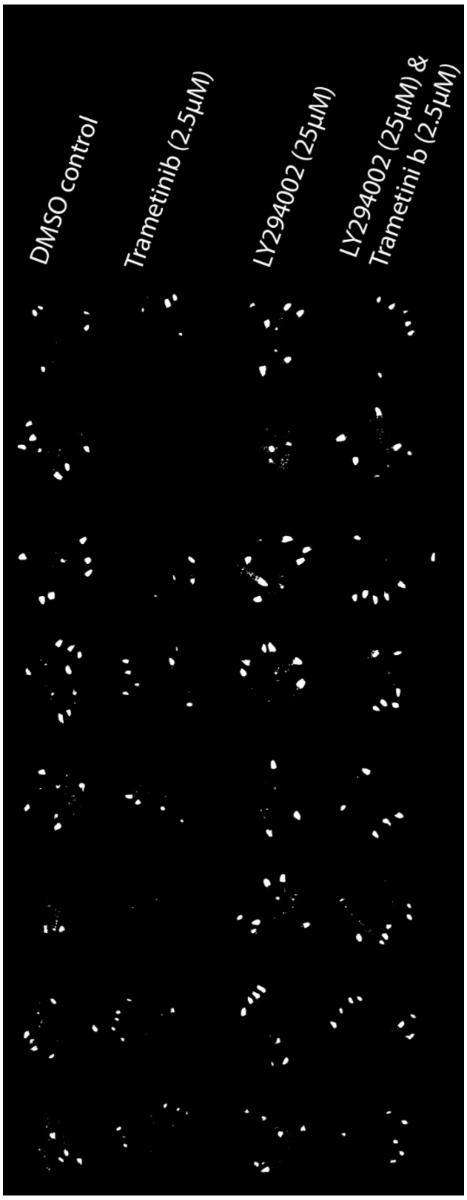


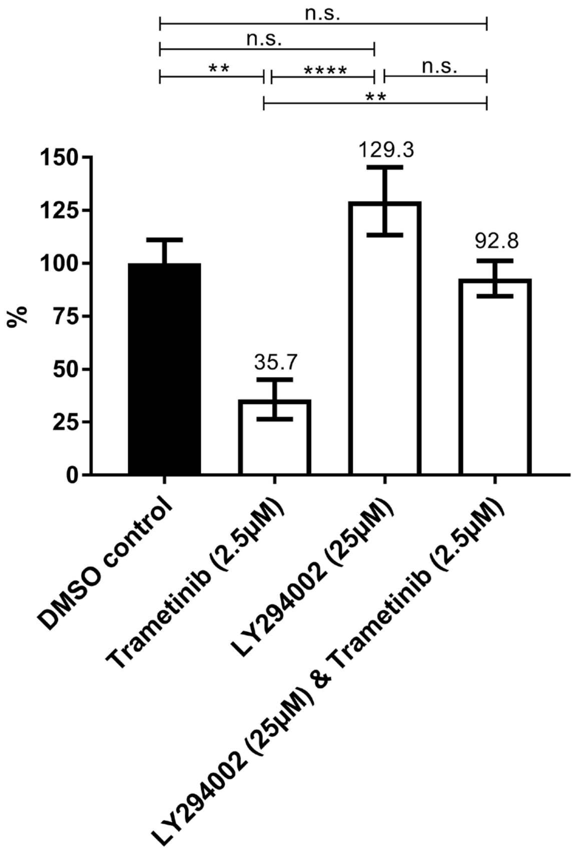


**Left** – Partial binarized image of screening plate showing various LY294002 treatments. **Right** – Mean GFP pixel area, represented as a percentage of mean DMSO pixel area, for various LY294002 treatments. N = 8 replicate wells per treatment. Error bars represent S.E.M. A one-way ANOVA, with a Tukey's multiple comparison test, was used to measure statistical significance. ** $p < 0.01$, **** $p < 0.0001$.

**Compound:** NU7441
**Reported Function:** Potent and selective DNA-PK inhibitor

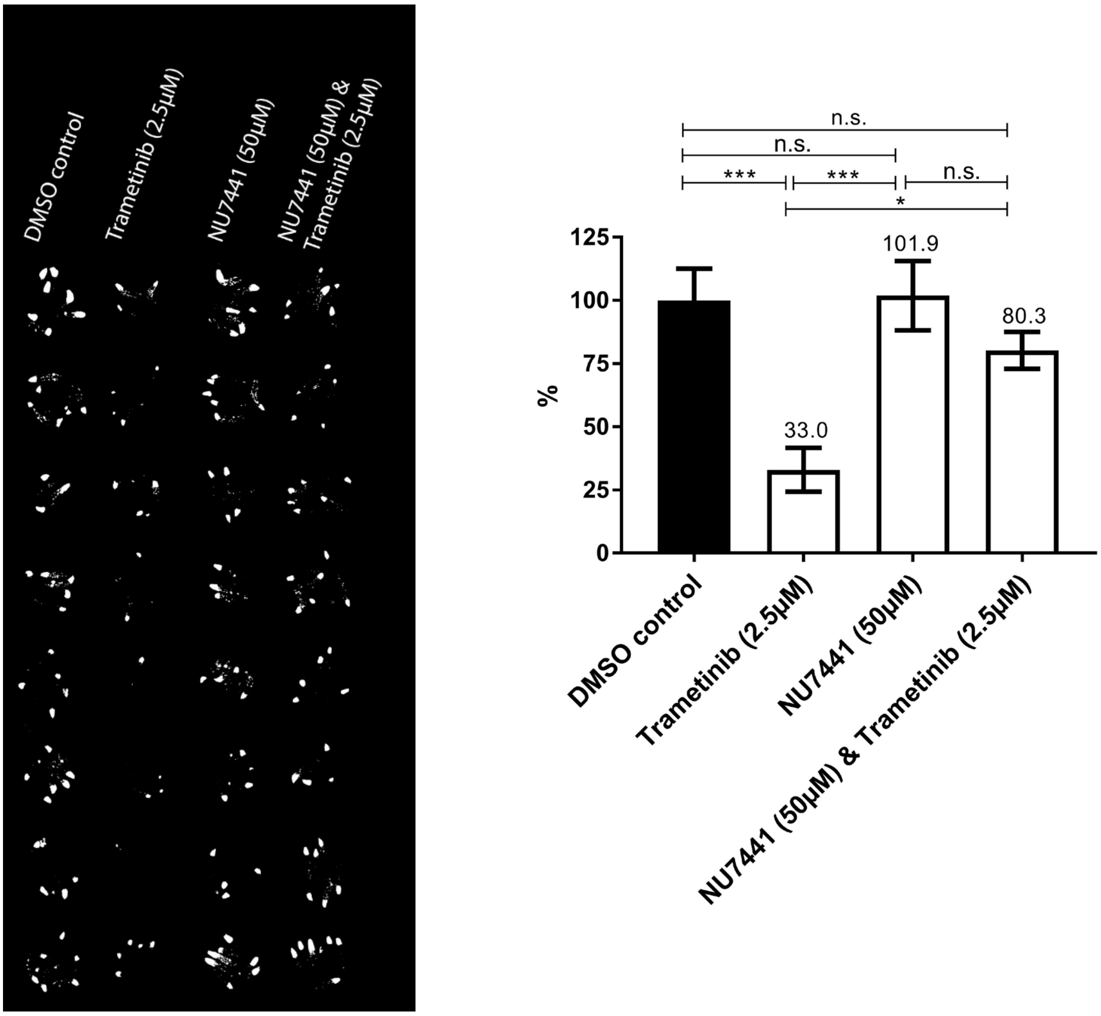


**Left** – Partial binarized image of screening plate showing various NU7441 treatments. **Right** – Mean GFP pixel area, represented as a percentage of mean DMSO pixel area, for various NU7441 treatments. N = 8 replicate wells per treatment. Error bars represent S.E.M. A one-way ANOVA, with a Tukey's multiple comparison test, was used to measure statistical significance. * $p < 0.05$, *** $p < 0.001$, **** $p < 0.0001$.

**Compound:** Propylpyrazole Triol
**Reported Function:** Estrogen receptor α agonist

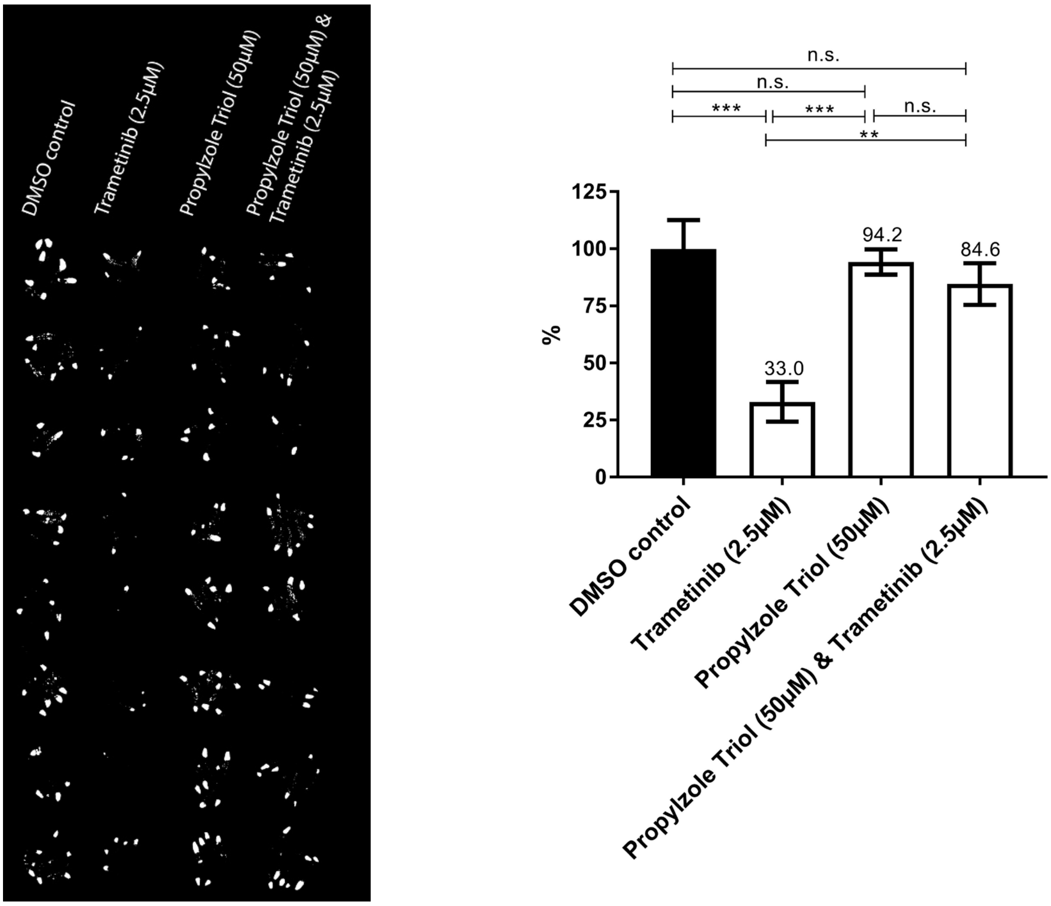


**Left** – Partial binarized image of screening plate showing various Propylpyrazole Triol treatments. **Right** Mean GFP pixel area, represented as a percentage of mean DMSO pixel area, for various Propylpyrazole Triol treatments. N = 8 replicate wells per treatment. Error bars represent S.E.M. A one-way ANOVA, with a Tukey's multiple comparison test, was used to measure statistical significance. ** $p < 0.01$, *** $p < 0.001$.

**Compound:** R1487 (hydrochloride)
**Reported Function:** Selective inhibitor of p38α

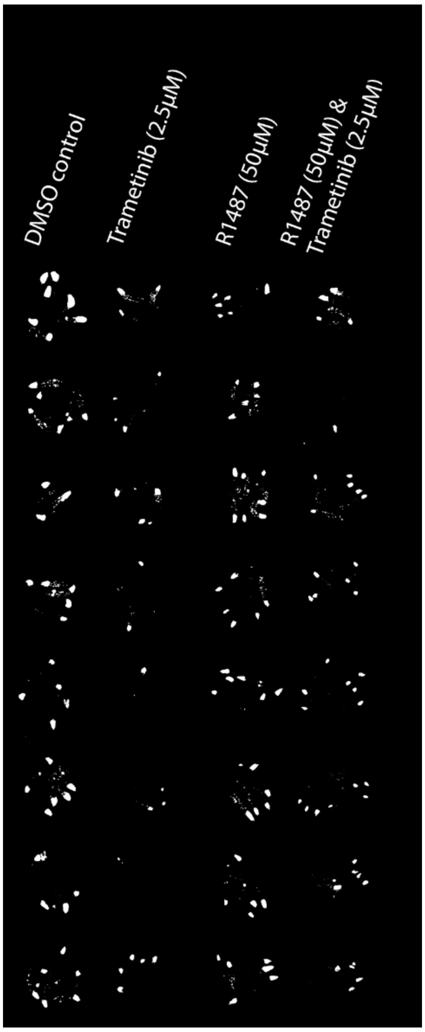


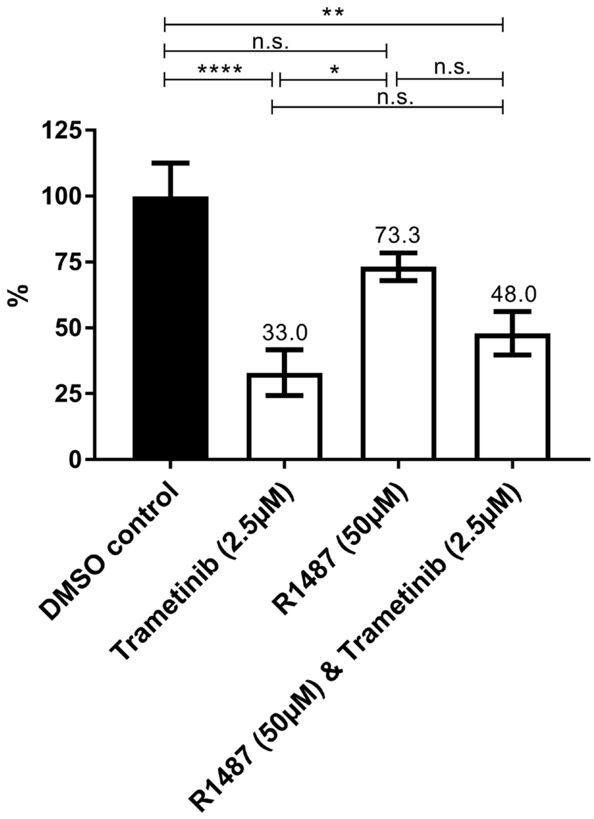


**Left** – Partial binarized image of screening plate showing various R1487treatments. **Right** – Mean GFP pixel area, represented as a percentage of mean DMSO pixel area, for various R1487 treatments. N = 8 replicate wells per treatment. Error bars represent S.E.M. A one-way ANOVA, with a Tukey's multiple comparison test, was used to measure statistical significance. * $p < 0.05$, ** $p < 0.01$, **** $p < 0.0001$.

**Compound:** SB202190
**Reported Function:** Inhibits p38 and p38β2

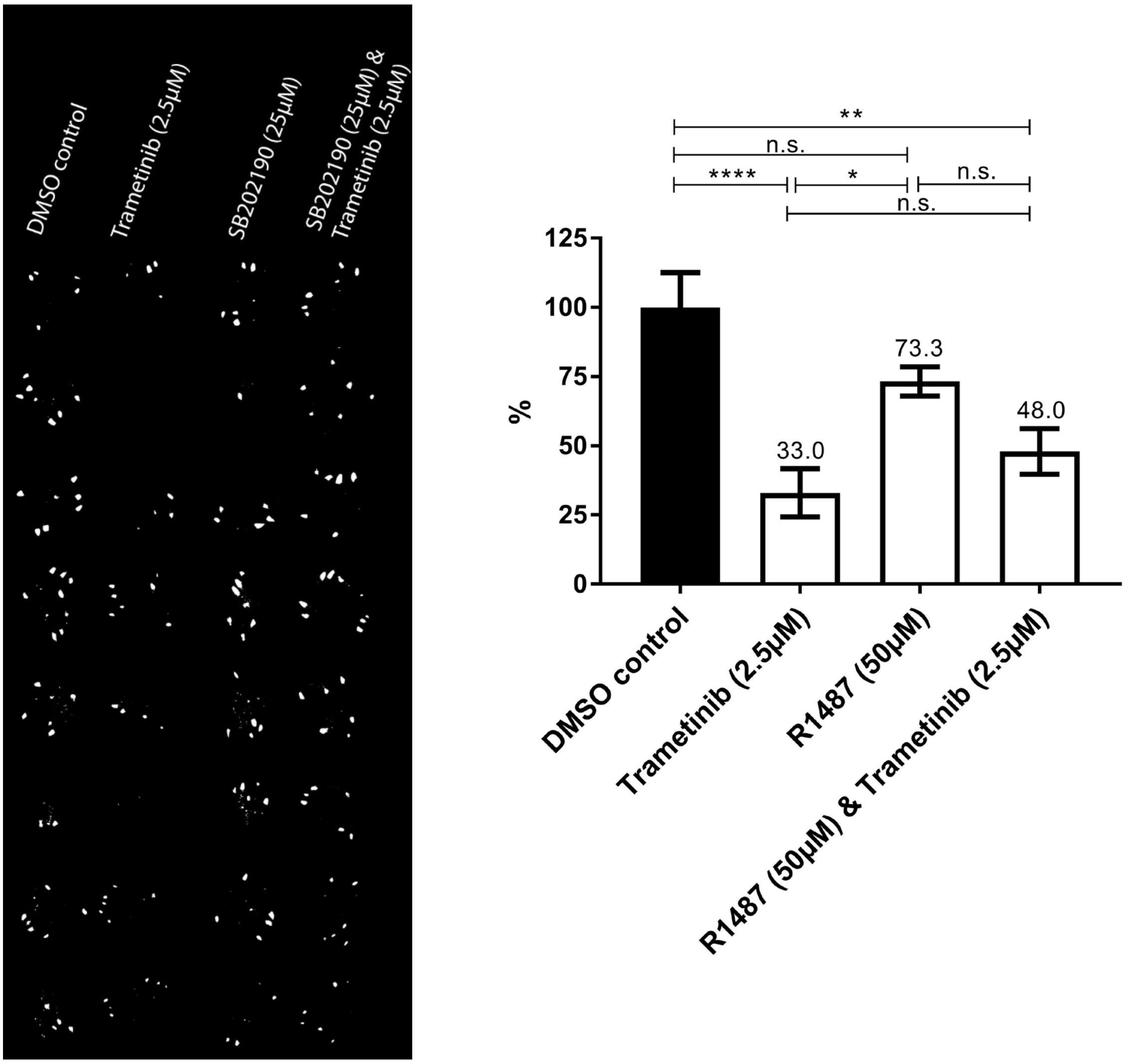


**Left** – Partial binarized image of screening plate showing various SB202190treatments. **Right** – Mean GFP pixel area, represented as a percentage of mean DMSO pixel area, for various SB202190treatments. N = 8 replicate wells per treatment. Error bars represent S.E.M. A one-way ANOVA, with a Tukey's multiple comparison test, was used to measure statistical significance. * $p < 0.05$, ** $p < 0.01$.

**Compound:** SKF81297C
**Reported Function:** A selective agonist of the dopamine D1-like receptor

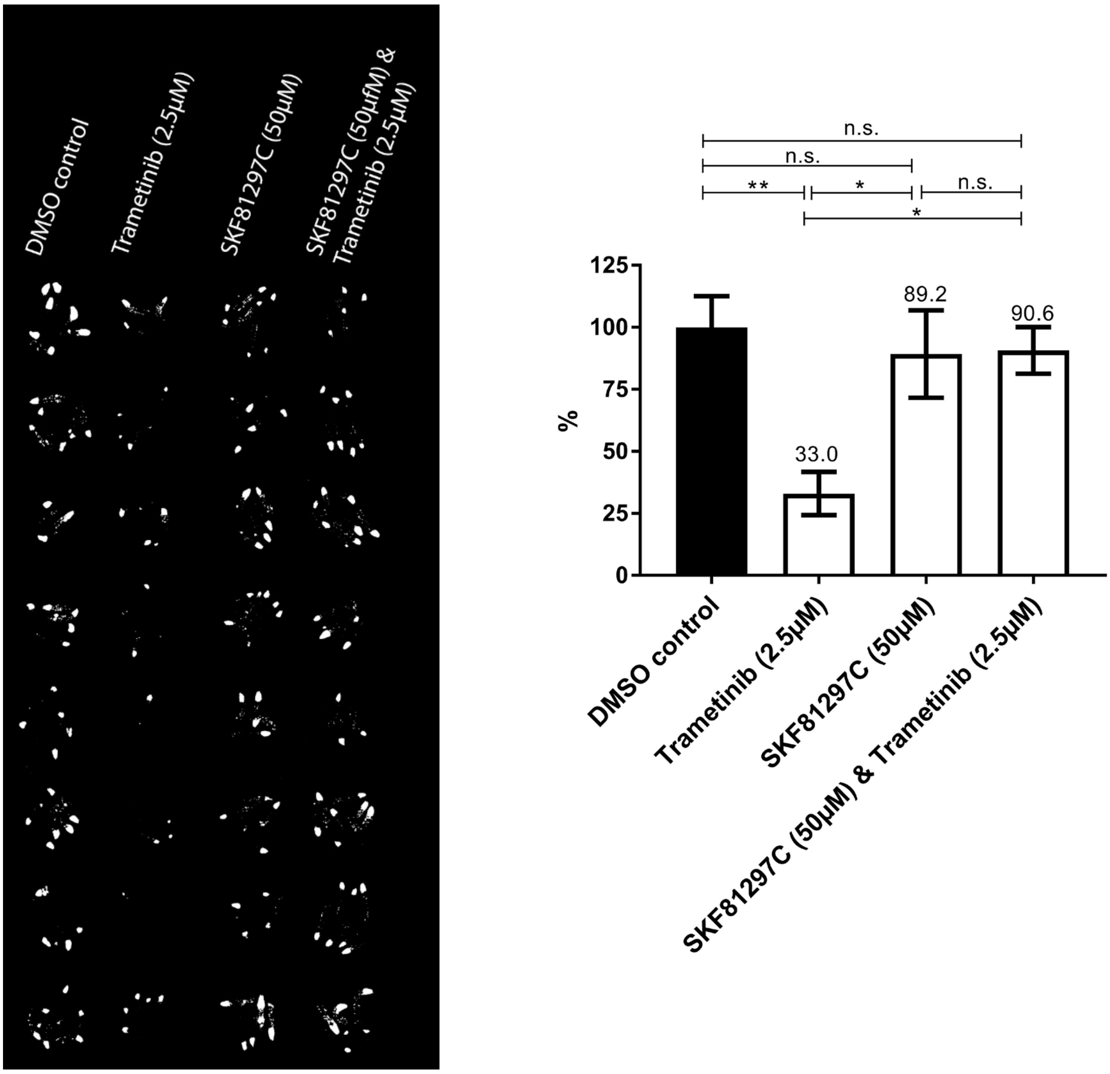


**Left** – Partial binarized image of screening plate showing various SKF81297C treatments. **Right** – Mean GFP pixel area, represented as a percentage of mean DMSO pixel area, for various SKF81297C treatments. N = 8 replicate wells per treatment. Error bars represent S.E.M. A one-way ANOVA, with a Tukey's multiple comparison test, was used to measure statistical significance. * $p < 0.05$, ** $p < 0.01$.

**Compound:** XL413 (hydrochloride)
**Reported Function:** Potent, selective and ATP competitive inhibitor of Cdc7. Also shows potency on CK2, PIM1.

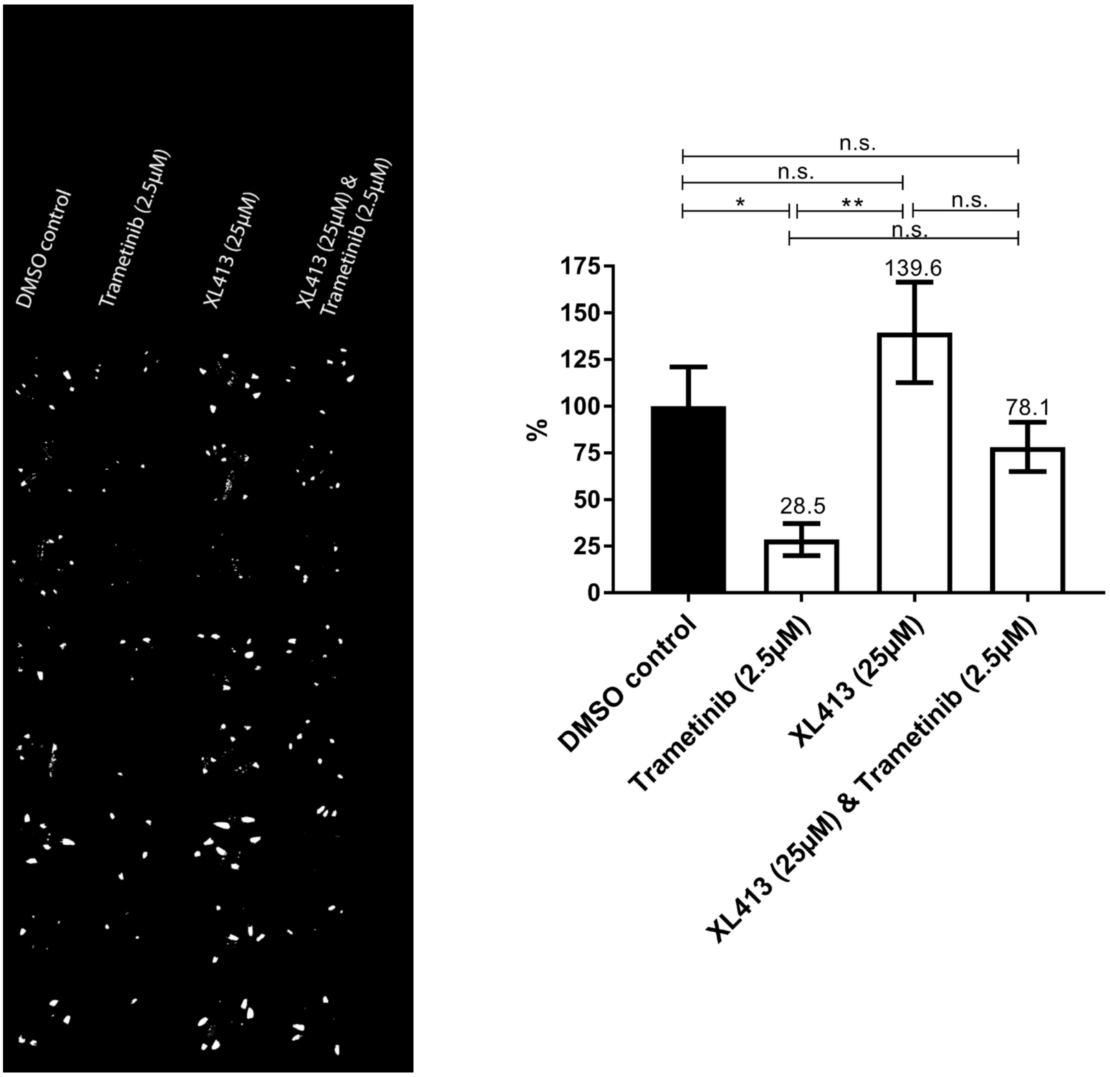


**Left** – Partial binarized image of screening plate showing various XL413treatments. **Right** – Mean GFP pixel area, represented as a percentage of mean DMSO pixel area, for various XL413treatments. N = 8 replicate wells per treatment. Error bars represent S.E.M. A one-way ANOVA, with a Tukey's multiple comparison test, was used to measure statistical significance. * $p < 0.05$, ** $p < 0.01$.
