## Supplementary material for "A *Drosophila in vivo* chemical screen reveals that combination drug treatment targeting MEK and DGKα mitigates Ras-driven polarity-impaired tumourigenesis": Supp File 4

**Supplementary File 4. Code used to perform automated analyses of binarized images of 96 well plates containing larvae treated with drugs of interest.**

```
// Designed for analyzing GFP of a thresholded (binarized) 96 well plate
// Whole plate image is stiched together from 2-well images
// set "my_width" to half the width, in pixels, of a 2-well image
// set "my_height" to the height of the 2-well image
// Data is read out as one well at a time along a row (12 columns per row, 8 rows)
// E.g. A1, A2, A3, A4 etc.


my_width = 624;
my_height = 648;
loc_x = 0;
loc_y = 0;

for (j = 0; j < 8; j++) {
	loc_x = 0;
	for (i = 0; i < 12; i++) {
		makeRectangle(loc_x, loc_y, my_width, my_height);
		run("Analyze Particles...", "size=2-Infinity exclude clear summarize");
		loc_x += my_width + 1;
	}
	loc_y += my_height + 1;
}
```
