## Supplementary material for "A *Drosophila in vivo* chemical screen reveals that combination drug treatment targeting MEK and DGKα mitigates Ras-driven polarity-impaired tumourigenesis": Supp Tables 1-3

**Supplementary Tables.**

**Supplementary Table 1. Fly Stocks used in this study.**

A list of stocks used throughout the project where the stock names indicate the genotype of the fly stock. VDRC is the Vienna Drosophila Resource Centre. BDSC is the Bloomington Drosophila Stock Centre.

| **Stock Name** | **Additional Names** | **Source** | **Catalogue Number** | **Additional Information** |
| --- | --- | --- | --- | --- |
| *eyFLP, UAS-GFP ;; tub-GAL4, FRT82B, tub-GAL80 / TM6B-RFP* | MARCM82B |  |  |  |
| *UAS-Ras85D^V12^ ; FRT82B, scrib^1^ / TM6B* |  |  |  |  |
| *eyFLP ;; Act>>GAL4, UAS-GFP / TM6B* | eyFLPout  EAG | Louise Cheng |  |  |
| *eyFLP ; UAS-Ras85D^V12^, UAS-dlg1 RNAi / CyO tub-GAL80 ; Act>>GAL4, UAS-GFP / TM6B* | EAGRD | Konrad Basler |  | REFERENCE Willecke et al., 2011 |
| *hsFLP ; Act>>GAL4* |  |  |  |  |
| *UAS-5-HT1A RNAi* | *5-HT1A^RNAi^* | VDRC | v106094 |  |
| *UAS-5-HT1B RNAi* | *5-HT1B^RNAi^* | BDSC | 27634 |  |
| *UAS-5-HT2A RNAi* | *5-HT2A^RNAi^* | BDSC | 31882 |  |
| *UAS-5-HT2B RNAi* | *5-HT2B^RNAi^* | BDSC | 60488 |  |
| *UAS-5-HT7 RNAi* | *5-HT7^RNAi^* | BDSC | v330023 |  |
| *UAS-Dgk RNAi* | *Dgk^RNAi^* | VDRC | v38239 |  |
| *UAS-rdgA RNAi* | *rdgA^RNAi^* | VDRC | v3024 |  |
| *UAS-Dgkε RNAi* | *Dgkε^RNAi^* | VDRC | v330361 |  |
| *UAS-luciferase RNAi* | *luciferase^RNAi^* | BDSC | 31603 | Used as a non-targeting RNAi control |
| *w^1118^* |  | BDSC | 3605 | Used as a wildtype |

**Supplementary Table 2. Primers used in qRT-PCR analysis.**

| **Gene** | **Forward Primer (5’-3’)** | **Reverse Primer (5’-3’)** |
| --- | --- | --- |
| *Gapdh2* | GCAAGCAAGCCGATAGATAAACA | CGTTGGCGCCCTTATCAATG |
| *Dgk* | CGGTGGCCATTCAGGACTTT | CGAAGTCCAGTCCTCACCTG |
| *Dgkε* | ATGGAGGTGTTCGGCATTGT | GGTCTCCTTGACTTGTAGCCTT |
| *5-HT1A* | TTGCCGTCGATCGTTACTGG | GCCGTCCAAACGCAAAAGAT |
| *5-HT1B* | GGAATCGCAGCATAAACGGC | ACGAATACGTTGCCTATGATGG |
| *5-HT2A* | CTTTTTCGCAGGTTGGGTGG | GTTGAAGTTGCGATTGCCGT |
| *5-HT2B* | GGAATAAAACACGTCGGCGG | CGAGGCGTGATTCTTGGAGT |
| *5-HT7* | CGAGAAGAAAGCCGATCCGA | GCAGTCGGTTTCGCAGACTA |
| *rdgA* | CATTCCATCGCCGGAGGTAA | GGAACATGTCCAGGCCCATT |
| *Rpl32* | CCAGTCGGATCGATATGCTAA | GTTCGATCCGTAACCGATGT |

**Supplementary Table 3. Primary antibodies used in western blots**.

| **Antigen** | **Host Species** | **Dilution** | **Source** |
| --- | --- | --- | --- |
| pS6 | Rb | 1:2000 | Cell Signaling Technology #4858 |
| S6 | Rb | 1:10000 | Cell Signaling Technology #2217 |
| pAkt | Rb | 1:1000 | Cell Signaling Technology #4060 |
| Akt | Rb | 1:1000 | Cell Signaling Technology #4691 |
| pERK | Ms | 1:1000 | Cell Signaling Technology #9106 |
| ERK | Rb | 1:500 | Cell Signaling Technology #9102 |
